# Dysregulation of transcription networks regulating oligodendrogenesis in age-related decline in CNS remyelination

**DOI:** 10.1101/2025.11.14.688494

**Authors:** Penelope Dimas, Samuel Morabito, Khalil S. Rawji, Qing Wang, Fynn N Krause, Juan Cubillos, Nejma Belaadi, Peggy Assinck, Zechuan Shi, Zhenkun Cao, Holger Heyn, Riki Kawaguchi, Daniel H. Geschwind, Vivek Swarup, Robin JM Franklin

## Abstract

In demyelinating diseases like multiple sclerosis (MS), efficient remyelination is critical for functional recovery. Remyelination efficiency declines with age, and is linked to progressive disability. The gene regulatory network underlying remyelination, and how it is altered with aging, remains unclear. Here we present a comparative single-nucleus RNA and ATAC sequencing analysis of remyelination in young and aged mice. We identified gene modules dynamically expressed throughout oligodendrocyte differentiation, revealing age-dependent changes in key processes related to myelination. Multi-omic analysis allowed us to map the regulatory network driving efficient remyelination within oligodendrocyte lineage cells in young mice. We highlight key transcription factors in the network dysregulated with age, and we describe similar dysregulations in MS lesions. Modifying the expression of these transcription factors in primary oligodendrocyte progenitor cells impacts differentiation. These findings provide a foundational understanding of this regenerative process in the context of aging and in chronic demyelinating diseases.

## Introduction

Remyelination in the central nervous system (CNS) is a spontaneous regenerative process following demyelination in which adult oligodendrocyte progenitor cells (OPCs) become activated, proliferate, and differentiate into new myelin-forming oligodendrocytes^1^. Newly generated myelin sheaths provide critical neuroprotective support by enabling the restoration of conduction velocity and the provision of metabolic support to the underlying axon^2^. As with most regenerative processes, the efficiency of remyelination declines with age^3^. This age-associated decline underlies the clinical progression of multiple sclerosis (MS) into its chronic and progressively degenerative stage^4^. Elucidating the mechanisms of remyelination and how these change with age is critical for understanding age-related remyelination failure and how to overcome it therapeutically^5^.

The process of remyelination is fundamentally controlled by transcription factors (TFs), which orchestrate the differentiation and maturation of OPCs into myelin-producing oligodendrocytes. While many TFs, including OLIG1, OLIG2, SOX10, MYRF, NKX2.2, and SOX2^6–11^ have been identified as crucial for both developmental and regenerative myelination, their roles have often been studied in isolation. We lack a comprehensive understanding of how these TFs work together in an interdependent network to drive lineage progression. Establishing a hierarchical network of TFs driving remyelination not only provides a deeper understanding of the molecular basis of remyelination but also a framework for addressing the hypothesis that age-related remyelination failure arises when this network goes awry.

Here we used the lysolecithin mouse model of demyelination-remyelination coupled with single nucleus RNA sequencing (snRNA-seq) and single-nucleus ATAC sequencing (snATAC-seq) at key stages in the remyelination process to generate a comprehensive cellular map of remyelination in young and old mice. We devised a unique computational strategy to leverage our multi-omic data. We first identified 10 dynamic gene modules that represent distinct biological functions throughout the oligodendrocyte differentiation trajectory and uncovered significant age-related changes in their activity. We then used this information to construct gene regulatory networks (GRNs) and to identify processive cascades of TF activation throughout oligodendrocyte maturation. Our analysis identified key TFs that were not only expressed differentially in the oligodendrocyte lineage of young versus aging mice but also in human MS lesions. Experimental manipulation of key TFs significantly altered OPC differentiation and activation of downstream targets, demonstrating their functional importance and the validity of these GRNs.

## Results

### snRNAseq and snATACseq reveal cellular and chromatin landscape of remyelination in the young and aged CNS

To characterize the transcriptional dynamics underlying remyelination, we induced focal demyelinated lesions in the ventrolateral spinal cords of young adult (2-3 months) and aging (14-15 months) mice. Tissue was collected for snRNA-seq and snATAC-seq (10X Genomics v3) from naïve mice and at days 5, 14, and 30 following demyelination (*n* = 2 per timepoint and age), corresponding to phases of OPC recruitment/proliferation, OPC differentiation and active myelination, and completed remyelination (in young mice) respectively^12^ (Fig. 1a). After quality control filtering, a total of 157,563 nuclei were retained for snRNA-seq and 52,688 nuclei for snATAC-seq (Supplementary Figs. 1-2; Supplementary Table 1). Multi-omic integration and unbiased clustering analysis^13–15^ revealed all major cell lineages of the CNS (Fig. 1b; Supplementary Figs. 3-4). As expected for a white matter (WM) enriched tissue, glial cells were the most abundant cell type, with the majority of the 116,932 glial nuclei belonging to the oligodendrocyte lineage.

**Fig. 1.**
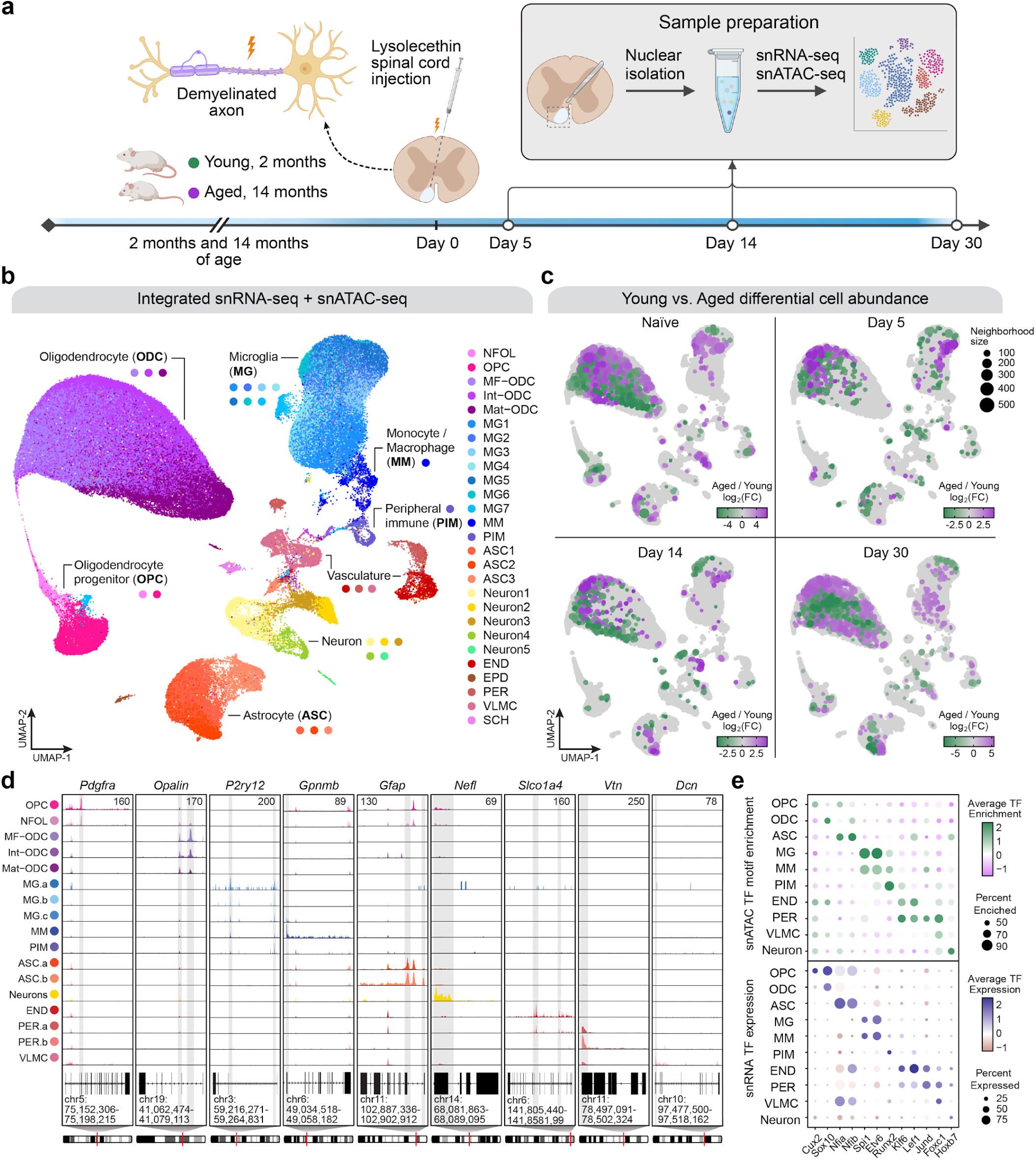
snATAC-seq and snRNA-seq of de- and remyelinating spinal cord lesions in young and aging mice. **a**, We induced focal demyelinated lesions in the ventrolateral spinal cords of young (2-3 months) and aged (14-15 months) mice, and sacrificed the mice at days 5, 14, and 30 post-demyelination, corresponding to the remyelination phases of OPC recruitment, OPC differentiation and completed remyelination (in young mice) respectively. snATAC-seq and snRNA-seq were performed in spinal cord tissue from these mice (n=2 per age and timepoint). **b**, Uniform manifold approximation and projection (UMAP) plot depicting a two-dimensional view of the integrated cellular neighborhood graph of 52,688 snATAC-seq profiles and 157,563 snRNA-seq profiles. Points represent cells, colored by cell-type annotations derived from unbiased clustering analysis. NFOL, newly formed oligodendrocyte; OPC, oligodendrocyte progenitor cell; MF-ODC, myelin-forming oligodendrocyte; Int-ODC, intermediate oligodendrocyte; Mat-ODC, mature oligodendrocyte; MG, microglia; MM, monocyte/macrophage; PIM, peripheral immune cell; ASC, astrocyte; END, endothelial cell; EPD, ependymal cell; PER, pericyte; VLMC, vascular leptomeningeal cell; SCH, Schwann cell. **c**, Differential cell abundance results comparing aged vs. young within the four experimental timepoints. Points in the UMAP represent cellular neighborhoods from Milo^16^, colored by the effect size of the comparison as log_2_(fold-change). **d**, Chromatin accessibility plots showing regions of accessibility (peaks) present in cell-type specific genes for each of the respective cell clusters. **e**, Dot plots displaying the average motif enrichment (snATAC-seq) and gene expression (snRNA-seq) for cell-type specific TFs for each of the cell clusters.

Differential cell abundance analysis^16^ comparing age groups at each experimental timepoint details the compositional shifts throughout remyelination (Fig. 1c, Supplementary Fig. 3d). This analysis revealed differentially enriched cell states with respect to aging across all cell lineages. Lesion samples showed a characteristic shift in their cellular composition enriched in immune cells, astrocytes, and OPCs in the early stages of remyelination in both young and aging mice. We identified cell-type specific patterns of gene expression and chromatin accessibility in the transcriptomic and epigenomic datasets (Fig. 1d, Supplementary Table 2). We also estimated genome-wide TF motif enrichment^17^ at single-nucleus resolution in the epigenomic dataset and found similarities between motif enrichment and TF expression (Fig. 1e). Furthermore, projecting cell-type labels from the snRNA-seq dataset to the snATAC-seq dataset via label transfer analysis demonstrates concordance across data modalities (Supplementary Fig. 4).

### Multi-omic trajectory analysis uncovers age-dependent dysregulation of dynamic gene co-expression networks

Simultaneous analysis of transcriptional changes based on gene expression and epigenetic changes based on chromatin accessibility allows for comprehensive analysis of TF regulatory activity. We first set out to understand the TF-driven, dynamic changes in gene expression that occur during oligodendrocyte differentiation and maturation during remyelination and how these may be perturbed with aging. To this end we first performed pseudotime trajectory analysis^18^ on oligodendrocyte lineage cells (OLCs, Fig. 2a). In both age groups, we could follow lineage progression from quiescent OPCs to proliferating OPCs, newly formed oligodendrocytes (NFOL), myelin-forming oligodendrocytes (MF-ODC), intermediate oligodendrocytes (Int-ODC), into fully mature oligodendrocytes (Mat-ODC) (Fig. 2a). We applied chromatin co-accessibility analysis^19^ in the OLC snATAC-seq dataset from young and aged mice to identify cis-regulatory linkages between promoters and putative enhancer elements (Fig. 2b). Using the snRNA-seq dataset, we identified 7,830 dynamically expressed genes across the oligodendrocyte trajectory (t-DEGs) in young and 7,245 in aging, with 6,168 genes with shared dynamic expression patterns in both age groups (Fig. 2c).

**Fig. 2.**
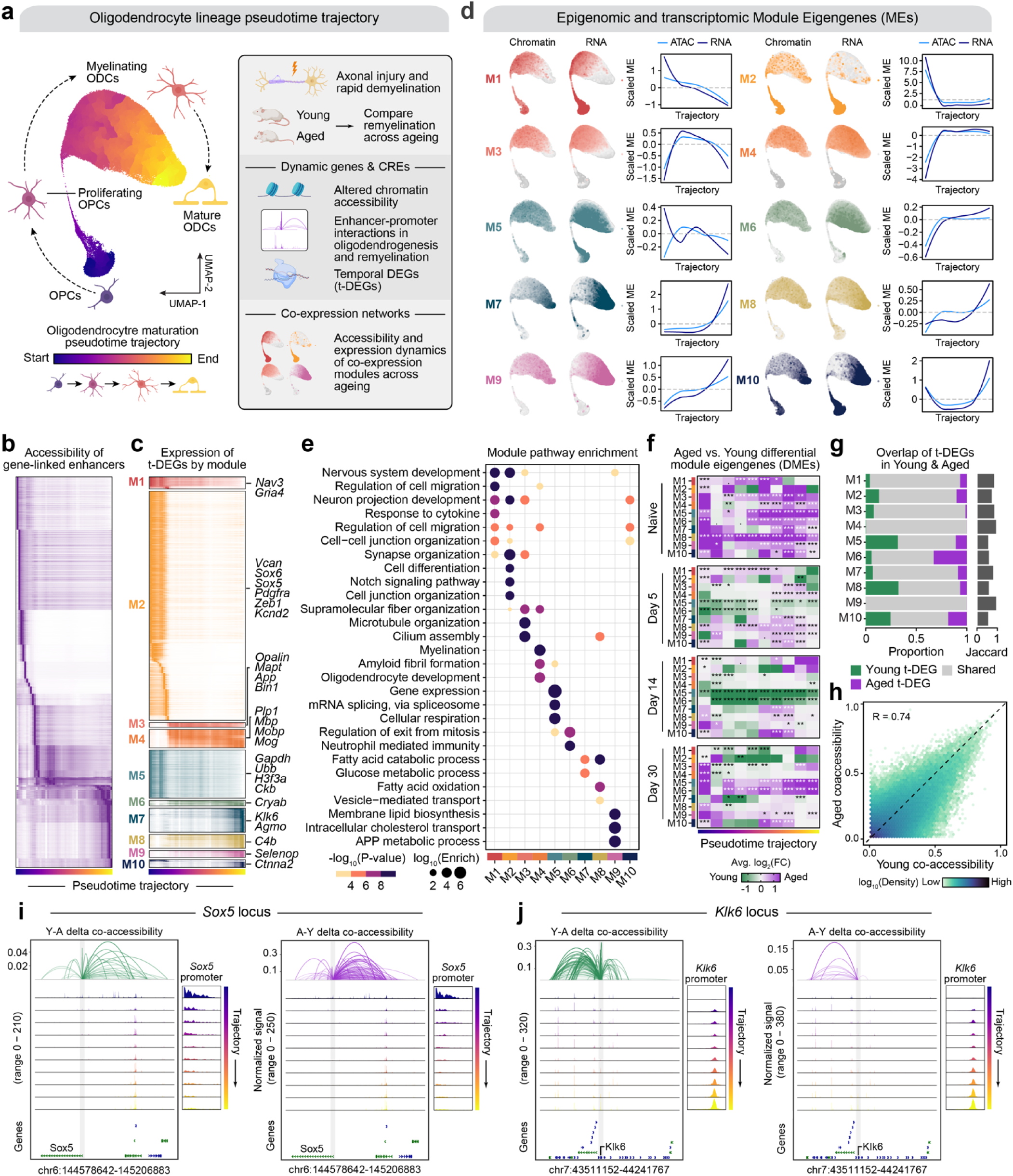
Trajectory analysis reveals dynamic signatures of oligodendrocyte maturation. **a**, Overview of the pseudotime trajectory analysis on the integrated snATAC-seq and snRNA-seq oligodendrocyte lineage data. A pseudotime trajectory was defined using Monocle3^18^. Trajectory dynamically expressed genes (t-DEGs) were identified in young and aged mice, and we performed co-expression network analysis of these genes using hdWGCNA^20^. We also linked genes to putative enhancers in young and aged mice via chromatin co-accessibility analysis. Illustrations were created using Biorender.com. **b**, Heatmap of scaled chromatin accessibility values for all gene-linked enhancer regions in 50 uniform bins across the pseudotime trajectory. **c**, Heatmap of scaled gene expression values for all oligodendrocyte t-DEGs in 50 uniform pseudotime bins. Genes are grouped by their co-expression module assignment, and ordered by pseudotime trajectory ranks. Selected module hub genes are annotated. **d**, Gene expression and chromatin accessibility dynamics of the ten oligodendrocyte lineage co-expression modules. Module eigengene (ME) UMAP feature plots of snATAC-seq and snRNA-seq, and ME trajectory line plots. **e**, GO term enrichment results for each co-expression module. **f**, Heatmaps showing DME results comparing aged vs. young for each module across the 10 uniform bins across the pseudotime trajectory, split by the four experimental timepoints. DME comparison was made using a two-sided Wilcoxon rank sum test with Bonferroni P-value adjustment. * P ≤ 0.05, ** P ≤ 0.01, *** P ≤ 0.001. **g**, Proportion of genes in each module that were young t-DEGs (left), aged t-DEGs (right), or shared (middle). The number of genes in each set are annotated. Jaccard index of the overlapping sets are shown in the bar plot on the right. H, Density heatmap showing distributions of chromatin co-accessibility values for promoter-enhancer pairs detected in the oligodendrocyte lineage in young vs. aged mice. Pearson correlation value and significance level are annotated. **i**-**j**, Epigenomic landscape of the *Sox5* locus (**i**) and the *Klk6* locus (**j**) in young (left) and aged (right) mice. Top: Delta chromatin co-accessibility (aged – young) showing enhancer links flanking the *Sox5* or *Klk6* promoter. Middle: Normalized chromatin coverage plots in 10 uniform bins across the pseudotime trajectory. Bottom: gene models at this genomic locus.

To characterize major gene expression programs driving remyelination in young and aging mice, we performed gene co-expression network analysis^20^ using young and aged t-DEGs combined (8,907 genes total), revealing 10 modules (M1 – M10) of highly co-expressed genes (Fig 2c-d, Supplementary Fig. 5, Supplementary Table 5). We calculated module eigengenes (MEs), representative gene expression profiles based on the entire group of genes present in each module, revealing dynamic expression patterns throughout the oligodendrocyte trajectory (Fig. 2d). Genes and pathways enriched in these modules highlight key regulatory relationships of gene networks underlying the biology of OLCs specification, differentiation and myelination (Fig. 2e). We projected these modules into the snATAC-seq dataset to assess the epigenomic activity of each module throughout the trajectory via epigenomic MEs (Fig. 2d), and we found a high degree of concordance across transcriptomic MEs and epigenomic MEs (median Pearson correlation per datasets: R = 0.949), suggesting tightly coordinated systemic changes that are conserved from chromatin accessibility to gene expression.

Modules M1 and M2 are expressed in the early stages in the oligodendrocyte lineage (OPC stage) and are associated with development-related Gene Ontology (GO) terms such as “regulation of cell migration”, “response to cytokine”, “cell differentiation”, and the “Notch signalling pathway” (Figs. 2d-e). Modules M3, M4, and M6 are expressed in the intermediate stages (NFOLs and MF-ODCs) and are associated with GO terms such as “myelination”, “oligodendrocyte development”, and “regulation of exit from mitosis” (Figs. 2d-e). Well known myelin-associated genes including *Plp1*, *Mbp*, *Mobp*, and *Mog* were all among the hub genes of module M4, further supporting its role in regulating myelination. Modules M7 – M9 are mostly enriched in late stages in the lineage (mature oligodendrocytes) and are associated with GO terms such as “membrane lipid biosynthesis” and “intracellular cholesterol transport” (Figs. 2d-e). Interestingly, M10 is activated both in progenitors and mature cells, indicating repression of the module in immature oligodendrocyte stages during the phase of active remyelination (Figs. 2d-e). When examining differentially expressed modules between young and aged OLCs, we found significant differences in most modules at all 4 time points (Naïve, Day 5, Day 14, and Day 30). Most notable was a downregulation in aged OLCs in Modules 5 and 6 at Days 5 and 14, suggesting a dysregulation in gene networks underlying gene-regulatory machinery like “mRNA splicing, via spliceosome”, and “regulation of exit from mitosis” (Fig. 2f). Additionally, we found significant differences in the myelination-related module M4 at key points along the trajectory, for example M4 was upregulated in aged OLCs at the end of the trajectory, suggesting a delayed timing in the expression programs related to myelination with respect to aging (Fig. 2f). We inspected the composition of the gene sets of each module with respect to the sets of t-DEGs identified in young and aging mice (Fig. 2g). In each module, most genes were dynamically expressed in both young and aging mice. Remarkably, 319 out of 320 genes in the myelination module M4 were t-DEGs among both age groups, while other modules have higher proportions of age-specific t-DEGs such as the Mat-ODC modules M6 and M8. Furthermore, we note that the largest module M2 is primarily expressed in OPCs (Figs. 2b-d), and we performed a subsequent network analysis of these genes in OPCs only, defining 18 OPC-specific sub-modules with distinct expression dynamics and associated biological functions (OPC-M1 – OPC-M18; Supplementary Fig. 6; Supplementary Table 6).

Next, we tested whether the age-related changes we observed in the oligodendrocyte lineage were due to a broad, global alteration in chromatin accessibility with age. By comparing the strength of promoter-enhancer co-accessibility links across the genome, we found a high degree of conservation between young and aging OLCs (Pearson correlation R = 0.74, Fig. 2h). This suggests that the observed age-related changes are not caused by widespread epigenetic remodeling but rather by more specific disruptions. We found several promoter-enhancer pairs with altered co-accessibility strength were highly interconnected hub genes, identified in our analysis of gene co-expression network modules. We focused on the cis-regulatory landscapes of two specific genomic loci with promoter-enhancer differences during aging to exemplify this: *Sox5* and *Klk6*, which are hub genes of module M2 (expressed in OPCs) and M7 (expressed in mature ODCs) respectively (Figs. 2i-j). The transcription factor *Sox5* is expressed in OPCs and was shown to be involved in regulating the specification and differentiation of OPCs into myelinating ODCs^21,22^. The *Sox5* promoter region was only accessible in the beginning of the trajectory in the OPCs, and overall, we found stronger and more numerous promoter-enhancer co-accessibility links in the aging OLCs compared to young. At the *Klk6* locus, a gene that encodes a serine protease that is exclusively expressed in mature ODCs, we found an increase of promoter-enhancer links in young ODCs relative to aging ODCs. *Klk6* has been implicated in MS pathogenesis^23–25^, and *Klk6* inhibitor studies suggest that *Klk6* may suppress myelination by limiting pro-myelinating signals^26,27^. In line with literature, both of these changes highlight age-associated changes that limit the capacity of OLCs to differentiate and form fully myelinating ODCs^28,29^.

### Age-dependent dysregulation of the gene regulatory network of remyelination

Cell fate decisions are controlled by tightly regulated molecular cascades, ensuring proper spatiotemporal expression of specific gene programs^30^. To model the cascade of TF-TF regulation required for oligodendrocyte lineage progression during remyelination, we employed a unique computational strategy leveraging the integrated ATAC and RNA data of the oligodendrocyte lineage trajectory in young and aging remyelinating OLCs (Fig. 3a). We constructed an initial GRN^31^ based on TFs with binding motifs in annotated promoter regions and gene-linked enhancer regions via co-accessibility analysis. Using XGBoost regression models, we refined the network by identifying TFs that were most predictive for the observed expression of each gene. This process yielded a directed GRN, with TFs linked to their respective sets of target genes, i.e. regulons (Supplementary Tables 7-8).

**Fig. 3.**
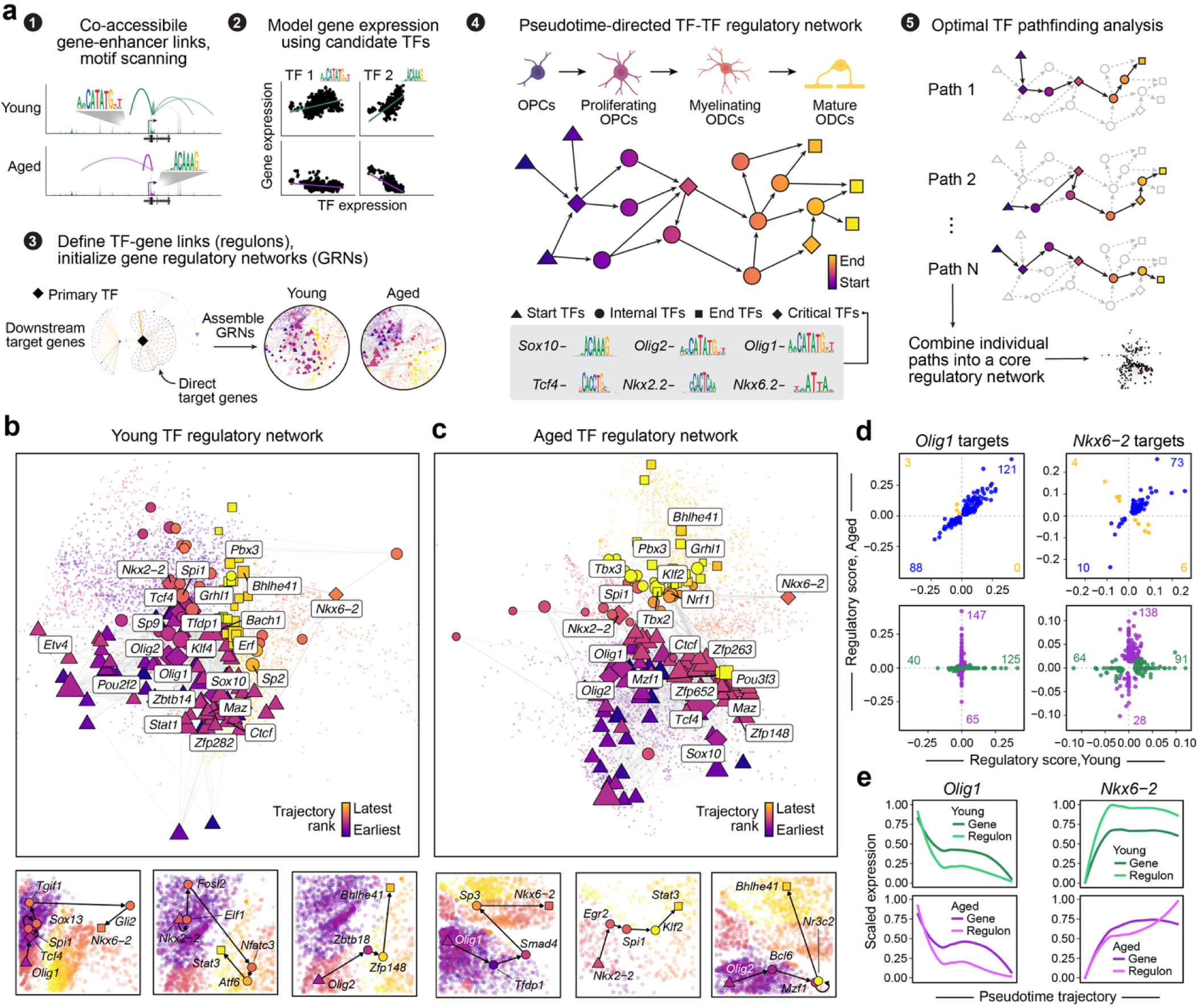
Transcription factor regulatory cascades underlying remyelination in aging mice. **a**, TF regulatory network analysis strategy. TF binding motif scanning in promoter regions and putative gene-linked enhancers for defining potential TF-gene relationships. XGBoost regression models the expression of each gene by potential regulators, ranking the top TFs per gene and defining the set of target genes per TF (regulons) comprising the GRN. We identified the most optimal paths in this directed network between TFs expressed in OPCs to TFs expressed in mature ODCs, revealing a core set of regulators in remyelination. Illustrations were created using <u>Biorender.com</u>. **b-c**, Subgraphs of the oligodendrocyte TF regulatory networks in young (b) and aged (c) mice showing nodes (TFs) and edges identified via optimal pathfinding analysis (other genes in the network shown in the background). Nodes are colored by pseudotime trajectory rank. Critical TFs (*Sox10*, *Olig2*, *Olig1*, *Tcf4*, *Nkx2.2*, and *Nkx6.2*) and most connected TFs in the network are labelled. Three TF cascades from optimal pathfinding analysis are shown below the main network: *Olig1* to *Nkx6.2*; *Nkx2.2* to *Stat3*; *Olig2* to *Bhlhe41*. **d**, Comparison of TF regulatory scores for the inferred regulons of *Olig1* and *Nkx6.2* in young vs. aged mice. Plots are faceted based on shared genes identified in the regulon in both young and aged (top) and unique genes (bottom). **e**, Scaled expression and aggregated TF regulon UCell^70^ signature scores along the pseudotime trajectory for *Olig1* and *Nkx6.2* in young and aged mice.

To pinpoint the precise sequence of TF activation, we used optimal pathfinding analysis (Fig. 3a). For optimal pathfinding we defined TFs that were highly expressed in OPCs as “start TFs”, and TFs expressed highly in Mat-ODCs as “end TFs”. We further stratified the network by ensuring that known key, “critical TFs” for oligodendrocyte lineage progression (such as *SOX10, OLIG2, OLIG1, TCF4,* MYRF, NKX2-2, and NKX6-2) were represented in the computed GRN, and assembled individual cascades into a core pseudotime-directed GRNs. This allowed us to map the regulatory cascades—the step-by-step TF activation events—that drive the maturation of OPCs into Mat-ODCs for young and aged OLC trajectories, which revealed distinct gene-regulatory patterns between the two age groups (Figs. 3b-c).

GRNs are highly complex systems with an intricate web of connections that provide both redundancy and resilience against minor perturbations. We hypothesized that for a perturbation to result in a severe pathology, such as that associated with aging, it must disrupt the function of TFs—or “hubs”—which possess a high degree of connectivity and control large numbers of downstream target genes. Our analysis revealed significant age-dependent changes in the connections and expression dynamics of key TF pairs, including OLIG1 – NKX6-2, NKX2-2 – STAT3, and OLIG2 – BHLHE41 (Figs. 3b-e; Supplementary Figs. 7-8). For instance, the TFs Olig1 and Nkx6-2 regulate distinct sets of genes in the young and aged GRNs, with over 100 unique target genes per age group (Fig. 3d). Furthermore, we found that the expression dynamics of NKX6-2 are temporally delayed in the aged GRN. This delay in the activation of its entire regulon (Fig. 3e) provides a compelling and plausible molecular mechanism that contributes to the well-documented age-dependent delay in oligodendrocyte differentiation and remyelination^29^.

### Identification of TFs that are linked to the age-dependent remyelination failure in MS

To determine if the transcriptional network changes we observed in aging mice are also relevant to human disease, we re-analyzed two published snRNA-seq datasets^32,33^ from human MS lesions and healthy WM (Fig. 4a; see Methods for details, Supplementary Fig. 9). We compared differentially expressed genes (DEGs) in MS lesions to those in young and aged OLCs during remyelination (Supplementary Tables 9-10; Supplementary Figs. 10-11). Given the known deficits in lineage progression of aging OLCs, we focused our analysis on DEGs that were also dynamically expressed along the OLC trajectory (t-DEGs) and were part of our previously computed regulatory network (Fig. 3). We further identified genes with a “switch-like” expression pattern—those with peak expression in a narrow window of the trajectory (Fig. 4b). We found 1,033 such switch-like genes that were shared between young and aged mice, a statistically significant overlap (Fig. 4c, one-sided Fisher’s exact test P = 0.00). These switch-like genes were most active during critical transitions in lineage progression, such as the differentiation from OPCs to immature oligodendrocytes. However, we found fewer of these genes active during these transitions in aging OLCs compared to young OLCs, again suggesting age-related differences in differentiation and maturation efficiency (Fig. 4c). Pathway enrichment analysis of these switch-like genes revealed enrichment of epigenetic processes and Wnt signaling exclusively in young OLCs, whereas those related to cellular metabolism and development were exclusively enriched in aging OLC switch-like genes (Fig. 4d-f).

**Fig. 4.**
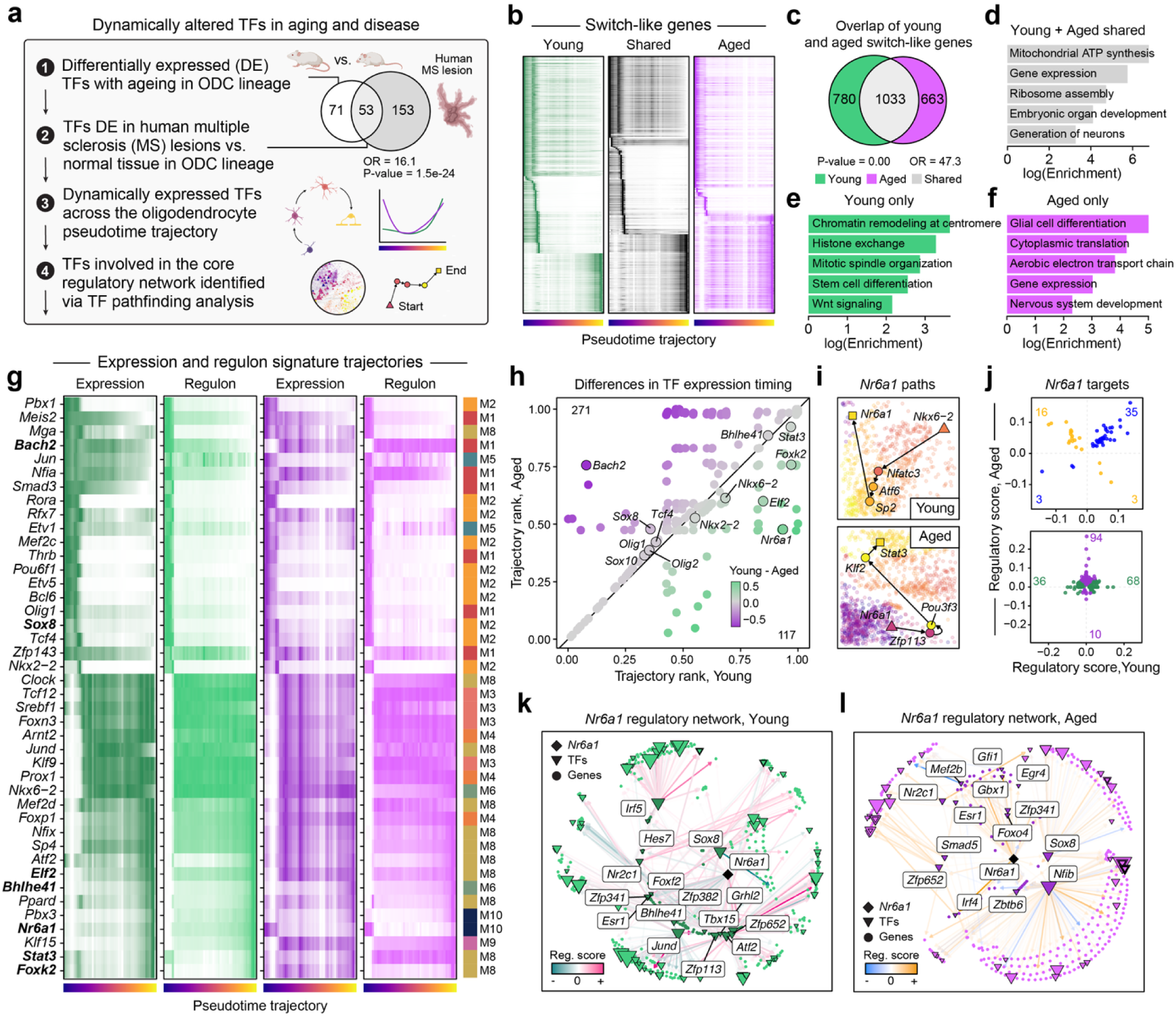
Differential regulation of key TFs in aging and MS. **a**, We define dynamic TFs dysregulated in aging and in human myelinating disease based on the following criteria: 1. TFs that are differentially expressed (DE) between young and aged mice in the oligodendrocyte lineage in our snRNA-seq dataset; 2. TFs that are DE in OLCs from MS lesions compared to healthy tissue in two public snRNA-seq datasets; 3. TFs that are dynamically expressed in the oligodendrocyte trajectory (t-DEGs); 4. TFs that represent a subgraph of the oligodendrocyte GRNs resulting from optimal pathfinding analysis. A set of 42 TFs meet these criteria. Illustrations were created using <u>Biorender.com</u>. **b**, Heatmaps of scaled gene expression values across 50 uniform pseudotime bins for genes with “switch-like” expression patterns in young and aged mice, and genes shared among both age groups. **c**, Euler diagram shows the overlap of switch-like genes identified in young and aged mice. One-sided Fisher’s exact test statistics are shown. OR: odds ratio. **d-f**, GO enrichment results for switch-like genes shared between young and aged (d), young only (e), and aged only (f). **g**, Heatmaps of scaled gene expression and aggregated TF regulon UCell^70^ signature scores for the 42 prioritized TFs from (a). TFs are ordered by pseudotime trajectory ranking in young mice. Co-expression module assignments for each TF are shown on the right. TFs selected for downstream validation experiments are highlighted with bold text. **h**, Comparison of pseudotime trajectory ranks of dynamically expressed TFs. Critical TFs and TFs used for downstream validation are labeled. Points are colored by the relative difference in TF expression timing based on trajectory ranks between young and aged. **i**, TF cascades from optimal pathfinding analysis for *Nkx6-2* to *Nr6a1* in the young network (top) and *Nr6a1* to *Stat3* in the aged network. **j**, Comparison of TF regulatory scores for the inferred regulons of *Nr6a1* in young vs. aged mice. Plots are faceted based on shared genes identified in the regulon in both young and aged (top) and unique genes (bottom). **k-l**, *Nr6a1* regulatory network in young (k) and aged (l) mice. Nodes represent genes, and edges represent TF-gene regulatory relationships. Primary (direct) target genes representing the regulon of *Nr6a1* are shown as darker nodes, and secondary target genes (i.e., the regulons of other TFs targeted by *Nr6a1*) are shown in a lighter color. Other TFs targeted by *Nr6a1* are labeled.

We identified a significant overlap between the TFs dysregulated in the aging mouse oligodendrocyte lineage and those in human MS lesions, with 53 differentially expressed TFs shared between the two datasets (Fig. 4a, one-sided Fisher’s exact test P = 1.5e-24). Further refinement to include only TFs found in the GRN optimal pathfinding analysis revealed 42 TFs that were dysregulated in both our aging mouse model and in human MS lesions. This age-associated dysregulation was not confined to individual TF expression levels; it also impacted their associated regulons in both the aging mouse and human MS lesions (Fig. 4g). Pseudotime trajectory ranks of each TF (where lower ranks indicate maximum expression of the TF at an earlier stage in the trajectory and higher ranks indicate expression at a later stage) highlights that expression timing of core TFs such as *Olig1, Olig2,* and *Sox10* remained stable, but the temporal expression of other TFs, like *Bach2* and *Nr6a1*, was altered with age (Fig. 4h). Interestingly, co-expression network analysis had identified *Nr6a1* as a member of module M10, which is the only module active in both OPCs and in mature ODCs, suggesting a potential shift in its role as a regulator of mature ODCs to OPCs with aging (Fig. 2c-d). This shift is further supported by our optimal pathfinding analysis, showing the role of *Nr6a1* (GCNF) as a start TF in aged mice and an end TF in young mice (Fig. 4i). Further examination of target gene sets of Nr6a1 revealed significant age-related differences. We found the set of genes regulated by GCNF in young OLCs to be largely distinct from the set it regulates in aged OLCs, which suggests that the regulatory function of GCNF changes significantly with age (Figs. 4j-l). These results highlight the dysregulation of the aging transcription factor network driving oligodendrocyte lineage progression.This dysregulation occurs on multiple levels, including, altered transcription factor expression, timing of expression, and target gene regulation, affecting critical hub genes during major transitions in OLC lineage progression.

### Functional validation of critical TF dysregulation in oligodendrocyte differentiation

We selected seven TFs for experimental validation based on the criteria described in our analysis: BHLHE41, BACH2, ELF2, FOXK2, GCNF, SOX8, and STAT33 (Supplementary Figs. 12-13). Of these, only SOX8 and STAT3 had been previously implicated in OLC biology^34–37^. ELF2 had been suggested to have a positive correlation with myelination in a snRNAseq study of oligodendrocytes in MS^38^. The remaining 4 TFs have not previously been described in the context of OLC biology, but have been studied in the context of regulating proliferation and differentiation in other cell types^39–43^.

Most of the short listed TFs were upregulated with aging, and we therefore investigated gain-of-function effects in neonatal, primary rodent OPCs utilizing lentiviral vectors. Under proliferation conditions (Supplementary Fig. 14a), TF overexpression had no effect on proliferation (Supplementary Figs. 14b,c and Supplementary Figs. 15a, b). Overexpression of GCNF, however, seemed to have a positive effect on OPC survival (Supplementary Fig. 14d and Supplementary Figs. 15c, d). As most of these TFs are expressed later in the oligodendrocyte trajectory, we tested the gain-of-function effect on differentiation of neonatal OPCs (Fig. 5a). Individual overexpression of all 7 TFs significantly enhanced OPC differentiation, reaching or exceeding levels similar to those seen with the pro-differentiating hormone T3 (Figs. 5b, c and Supplementary Figs. 15e, f). These results corroborate previous reports on the pro-differentiation roles of these STAT3 and SOX8. However, this is the first observation to our knowledge of the other 5 TFs enhancing OPC differentiation. To further functionally validate the shortlisted TFs, we explored the effect of overexpression of these TFs on their respective downstream regulatory network. To this end we performed bulk RNA sequencing on the TF overexpressing OLCs. All TF with the exception of GCNF initially predicted to be involved in differentiation and maturation of OLC, therefore samples were collected and analyzed 24h after inducing differentiation through mitogen withdrawal, whereas GCNF overexpression samples were collected whilst in proliferation condition.

**Fig. 5.**
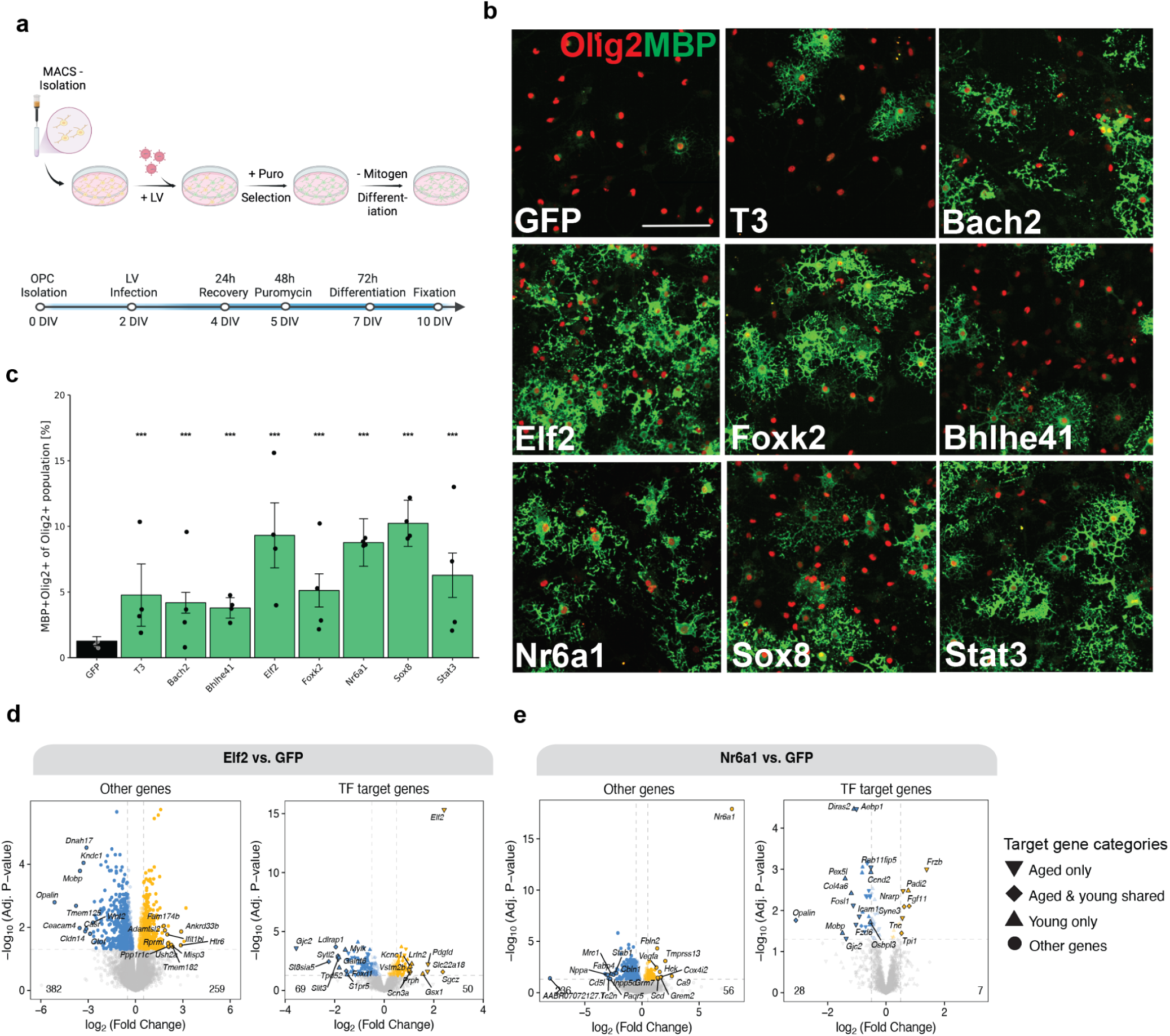
TF overexpression affects OPC differentiation and downstream gene expression. **a.** Diagram showing the experimental timeline of the OPC differentiation assay with lentiviral constructs for select transcription factor overexpression. Illustrations were created using <u>Biorender.com</u>. **b.** Representative images of control OPCs (GFP), control (with T3) OPCs, and OPCs transfected with the 7 select transcription factors in differentiation conditions and stained for Olig2 and myelin basic protein (MBP). **c.** Quantification of the percentage of differentiated OPCs (Olig2+MBP+) of total OLCs (Olig2+) in response to transcription factor overexpression treatments. Each data point shows the mean percentage across technical replicates (n=3) per biological replicate (n=4). Each bar shows the mean percentage across biological replicates (n=4), with propagated standard deviations indicated as error bars. Statistical analysis was performed using a linear mixed-effects model on the logit-transformed percentages, with transcription factor overexpression as a fixed effect and animal as a random effect. Green bars highlight transcription factor overexpression treatments with a significant effect on the percentage of differentiated OPCs compared to the control group (*** FDR-adjusted q < 0.001). Scale bar = 100 µm. **d.** Volcano plots showing differential gene expression results comparing transcription factor (TF) overexpression and GFP controls with bulk RNA-seq (n=4). The following TFs were overexpressed: *Elf2* (**d**) and *Nr6a1* (**e**). For each panel, the plot on the right shows primary & secondary target genes from both young and aged mice for the TF of interest, and the plot on the left shows all other genes.

Overexpression of all of these TFs drastically changed the transcriptional landscape of oligodendrocyte cells, including several TFs that are primary or secondary downstream of the overexpressed TF (Figs. 5d,e, Supplementary Fig. 16, Supplementary Table 11). Overall, these results confirm the validity of our computational analysis, and provide functional validation for the relevance of these TFs for remyelinating OLCs, and that their dysregulation is likely to contribute to the reduced ability of these cells for regeneration.

## Discussion

Our study provides a comprehensive, single-nucleus multi-omic dataset of remyelination, revealing the gene-regulatory dynamics orchestrating this vital regenerative process, and highlighting how it is fundamentally compromised with age. We identify key oligodendrocyte gene expression modules and regulatory networks, and characterize their dynamics throughout remyelination with respect to aging. Age-related shifts in the regulatory dynamics of these modules and networks coincide with the decline of remyelination efficiency seen throughout aging and in demyelinating diseases like MS^3^.

The high-resolution, integrated single-nucleus RNA and ATAC sequencing data allowed us to map the pseudotime trajectory of the oligodendrocyte lineage in unprecedented detail. We found that the core gene co-expression modules of the oligodendrocyte lineage, and many of the trajectory-dependent genes (t-DEGs), are shared between young and aged mice during remyelination. This suggests that the fundamental epigenomic and transcriptomic programs for remyelination are largely conserved with aging. However, a deeper analysis of the pseudotemporal dynamics in specific co-expression modules revealed significant age-dependent dysregulation, particularly among modules associated with myelination and oligodendrocyte development. This is exemplified by the downregulation of modules M5 and M6 in aged mice (naïve and day 30 timepoints), a finding that correlates with the well-established delay in oligodendrocyte differentiation and remyelination in older animals^3,29,44^.

Overall, we found age-related changes not be due to a broad change in chromatin accessibility, as we observed a high degree of conservation in promoter-enhancer co-accessibility. Instead, we found specific, localized disruptions in the cis-regulatory landscapes of key genes, such as Sox5 and Klk6. These region specific disruptions suggest that subtle age and region specific changes may underlie the profound failure in the precise gene regulatory control needed for successful maturation of OLCs. Our computational strategy allowed us to move beyond simple co-expression relationships towards modeling the directed, hierarchical regulatory cascades that drive the regulatory dynamics of oligodendrocyte lineage progression. We demonstrated that the age-related delay in remyelination is reflected in perturbations within this GRN. While core TFs like OLIG1, OLIG2, and SOX10 maintain their expression timing, the activity and regulatory influence of other crucial TFs are profoundly altered. A prime example is Nkx6-2, which shows a significant temporal delay in its regulatory signature in aged mice. This finding hints at a direct molecular explanation for the observed delayed differentiation, linking a specific TF-driven event to the macro-phenotype of remyelination failure.

By comparing our results with two published snRNA-seq datasets of human MS^32,33^, we establish a direct link between the transcriptional dysregulation in the aging mouse model of remyelination and the progressive pathology in human MS lesions. We identified a set of 42 conserved TFs that were i) differentially expressed between young and aged mice, ii) differentially expressed between MS lesions and healthy WM, iii) dynamically expressed across the OLCs (t-DEGs), and iv) implicated in the core GRN from TF pathfinding analysis. Ultimately, we further narrowed this to a group of 7 TFs for *in vitro* experimentation. The finding that these TFs are part of the core regulatory network and that their expression and regulatory function—such as the role of *Nr6a1* (GCNF) in young versus aged mice—are altered, provides compelling evidence that the similar GRN disruptions may contribute to age-related remyelination failure both in rodents and in human MS.

We could further confirm that this computed dysregulation in the aged GRN is of functional consequence to OLCs, as overexpression of each of the 7 TFs triggered changes in the ability of OPCs to form differentiated oligodendrocytes. Furthermore, experimental overexpression resulted in various changes in the downstream transcription factor network. Apart from SOX8 and STAT3, this is the first functional description of the other 5 TFs in OPCs. The pro-differentiation effects of these 7 TFs, together with the dysregulation in the aged GRN, further suggests that these key TFs may underlie the differentiation block observed in demyelinated lesions from aging rodents and chronic MS^29,45^.

Collectively, our work provides a foundational understanding of the GRN driving remyelination and identifies critical nodes of dysregulation within the GRN of the aging oligodendrocyte lineage. This work provides a foundational understanding of remyelination’s gene regulatory control and offers a powerful new framework for developing a network-based understanding of gene regulation during tissue regeneration and its age-dependent failure. Future work is needed to fully harness the power of this analysis and develop network modulating strategies through targeting single or combinations of critical nodal factors that are able to shift the entire network into a more youthful state.

## Methods

### Mice

All animal experiments were conducted in accordance with the United Kingdom Home Office regulations. Female C57Bl/6 mice from Charles River were used for lysolecithin demyelination at 2-3 months and 14-15 months of age and were housed under standard laboratory conditions on a 12-hour light/dark cycle. Mice were housed in groups of up to 5 animals without restriction to food and water.

### Lysolecithin-induced demyelination

Mice were anaesthetized with 2.5% isoflurane and administered buprenorphine (0.03 mg/kg, s.c.) as an analgesic. The thoracic spinal cord was exposed, after which 1.0 μL of 1.0% lysolecithin (L1381, Sigma) was injected to induce a focal demyelinating lesion into the ventrolateral funiculus over 2 minutes^46^. After injection, the needle was maintained in position for an additional 2 minutes, after which the mice were re-sutured and placed in a recovery chamber. Mice were monitored daily until time of sacrifice on Days 5, 14, and 30 after injection.

### Nuclei Isolation

Mice were perfused with RNase-free PBS under terminal anesthesia. A 6 mm segment of the spinal cord ventral hemisection, encompassing the lysolecithin lesion site, was dissected and snap-frozen on dry ice. The fresh frozen tissue was minced and homogenized in Nuclei EZ Lysis Buffer using a Dounce grinder (NUC101-1KT, Sigma Aldrich). The tissue homogenate was filtered and centrifuged through an OptiPrep™ Density Gradient Medium (Sigma Aldrich) to purify nuclei. Purified nuclei were stained with Trypan Blue and counted using a hemocytometer. The same nucleus suspension from each sample was used for both single-nucleus RNA-seq and single-nucleus ATAC-seq preparations.

### Single-nucleus RNA-seq and ATAC-seq Library Preparation and Sequencing

Single-nucleus RNA-seq libraries were prepared using the Chromium Next GEM Single Cell 3’ v3.1 Reagents Kit, and single-nucleus ATAC-seq libraries were prepared using the Chromium Next GEM Single Cell ATAC Reagents v1.1 Kit (10x Genomics), following the manufacturer’s instructions. Library quality was assessed with the TapeStation (Agilent) and quantified with Qubit (Thermo Fisher). Libraries were then pooled and sequenced on the NovaSeq 6000 platform (Illumina).

### Processing and initial analysis of the snATAC-seq dataset

For the snATAC-seq dataset, we aligned sequencing reads from each individual sample to the mouse reference genome (mm10) using cellranger-atac count (v 1.2.0). From cellranger-atac, we obtained a .bam file of mapped sequencing reads and a fragment file containing the genomic coordinates of each sequenced read-pair associated with cell barcodes, indicating a region of accessible chromatin in a single nucleus.

We imported these fragments files into ArchR^46^ for several downstream data processing tasks. ArchR initially quantifies chromatin accessibility in single nuclei by summarizing fragments in evenly spaced 500 bp genomic bins, referred to as a “tile matrix”. Additionally, ArchR uses this tile matrix to quantify a “gene score”, an epigenomic proxy of gene expression, using a model that summarizes the signal at a gene’s transcription start site (TSS) with a decaying weight throughout the gene body. Constructing the tile matrix and the gene score matrix was done using the ArchR function createArrowFiles, which also computes cell-level quality information like TSS enrichment. As an initial filter, we excluded low-quality nuclei with TSS enrichment lower than 4, or fewer than 1000 unique fragments. We also computed doublet scores using the addDoubletScores function and filtered out the inferred doublets (two or more distinct nuclei with the same barcode) from the downstream analysis.

We next sought to group cells together by their epigenomic signatures using dimensionality reduction and clustering analysis. We first performed iterative latent semantic indexing (iLSI) using the ArchR function addIterativeLSI, thereby producing a low-dimensional representation of the dataset. Harmony^47^ batch correction was applied to the iterative LSI matrix based on the sequencing batch. Using this batch-corrected representation, we then performed graph-based clustering with the Louvain algorithm using the function addClusters using a resolution of 0.8. We constructed a two-dimensional projection of the dataset by applying UMAP to the cell neighborhood graph previously used for clustering. Cell type identity labels were given to each cluster by inspecting gene score signatures and chromatin accessibility coverage tracks in each cluster for a panel of canonical cell type marker genes for cell types known to be found in the mouse spinal cord. We removed four clusters that showed conflicting marker gene scores, indicating additional doublet populations that were not previously identified. We next used MACS2 for peak calling in pseudo-bulk replicates for each of the snATAC-seq clusters. Peaks from these different analyses are then merged into a unified set of peaks using the iterative overlap approach in ArchR with the function addReproduciblePeakSet. Within this peakset, we tested for transcription motif enrichment in single nuclei using the chromVAR^17^ algorithm with the JASPAR2020^48^ motif database. After fully processing the dataset as described, we converted the ArchRProject object to SeuratObject format^49^ to facilitate downstream integration with the snRNA-seq dataset using Seurat (v 4.3)^50^ and Signac (v 1.10)^51^.

### Processing and initial analysis of the snRNA-seq dataset

For the snRNA-seq dataset, we aligned sequencing reads from each individual sample to the mouse reference transcriptome (mm10) using cellranger count (v 6.0). From cellranger count, we obtained a .bam file of mapped sequencing reads and a gene by barcode expression matrix (.mtx file) containing the gene number of UMI attributed to each gene for each barcode. We next used these gene expression matrices to identify potential doublets with the Scrublet^52^ algorithm, and we removed predicted doublets from our subsequent analysis.

For downstream data analysis with Seurat, we created a SeuratObject containing the merged gene expression matrices from the different samples. We removed low-quality nuclei based on the following thresholds: greater than 5% mitochondrial reads, fewer than 500 RNA counts, fewer than 200 genes detected, and greater than 5,000 genes detected. We performed dimensionality reduction and batch correction using online integrative non-negative matrix factorization (iNMF) with the R package Liger^53^. We used this iNMF data representation to compute a cell neighborhood graph with the FindNeighbors function, and to perform Louvain clustering analysis with the FindClusters function (resolution 0.5). We computed a two-dimensional projection of the snRNA-seq dataset by applying UMAP to the cell neighborhood graph. Cell type identity labels were given to each cluster by inspecting gene expression signatures in each cluster for a panel of canonical cell type marker genes for cell types known to be found in the mouse spinal cord. We employed a “one-versus-rest” strategy to perform differential gene expression analysis with MAST^54^ between a given cluster with all other clusters, enabling us to identify marker genes that are expressed specifically in each cluster.

### Integration of snATAC-seq and snRNA-seq datasets

We integrated our snATAC-seq and snRNA-seq datasets using Seurat to facilitate numerous downstream multi-omic analyses. Since we have the snATAC-seq gene score matrix, an epigenomic proxy of gene expression signal, we have a shared feature space to integrate the two datasets despite being measurements of distinct genomic features. In general, we use the snRNA-seq dataset as a reference in these integrative analyses because snATAC-seq data tends to be noisier than snRNA-seq data.

We first sought to perform a supervised transfer learning analysis to predict cell-type labels in our snATAC-seq dataset based on our snRNA-seq dataset. We used the Seurat function FindTransferAnchors to identify transfer anchors (groups of cells with highly similar molecular states based on their relationships in a nearest neighbor graph) using snRNA-seq as the reference and snATAC-seq as the query with canonical correlation analysis (CCA) as the underlying dimensionality reduction. We next used the Seurat function TransferData to compute a matrix of snATAC-seq nuclei by snRNA-seq clusters with each value in this matrix being the predicted probability of a snATAC-eq nuclei assigned to one of the snRNA-seq clusters. We then used the same anchors to obtain an imputed gene expression matrix for the snATAC-seq dataset using the Seurat TransferData function. We then merged the two datasets into one SeuratObject and computed an integrated UMAP of snRNA-seq and snATAC-seq.

### Linking genes to candidate cis-regulatory elements

Using our multi-omic dataset, we sought to identify regulatory links genes between candidate cis-regulatory elements (cCREs). This analysis is similar to our strategy for linking genes with cCREs in previous single-nucleus multi-omic study^13^. We first sought to identify pairs of chromatin accessibility peaks which were co-accessible in each cell type in our young and aged mice. This analysis was performed using the R package Cicero^19^, which identifies pairs of co-accessible peaks by correlating chromatin accessibility signals within aggregated metacell representations of the input data, penalizing the correlation based on genomic distance with a maximum allowable distance of 500kb. We performed this analysis separately for different cell types and experimental groups since these potential epigenetic regulatory relationships may be unique to certain biological conditions. While this analysis does not specifically focus on genes, to specifically inspect potential links between genes and cCREs, we first subset the set of co-accessible pairs identified by cicero where one peak flanks a gene’s promoter region and the other peak^55^ lies in a distal genomic region. For downstream analysis, we retained the top 20% peak-gene links by their co-accessibility scores.

### Multi-omic pseudotime trajectory analysis of the oligodendrocyte lineage

To thoroughly characterize the molecular signatures of OLCs in our dataset, we sought to perform an integrated multi-omic trajectory analysis using the R package Monocle3^18^ (v 1.3.1). For this analysis, we included all OLCs from both young and aged animals and all time points to be able to compare different groups along a unified trajectory. Using our integrated dataset as previously described, we first computed a principal graph in low-dimensional space using the Monocle3 function learn_graph with default parameters. We then ordered the cells along a pseudotime trajectory with the Monocle3 function order_cells, indicating the OPC cluster as the starting point.

We next sought to identify features that were changing along the trajectory. We used a generalized linear mixed effects model to perform differential gene expression analysis with respect to the pseudotime ordering as a continuous variable using the Monocle3 function fit_models. We also accounted for sequencing batch, number of UMI, and mitochondrial read content as model covariates. Trajectory dynamically expressed genes (t-DEGs) were denoted as those with a significant (Bonferroni adjusted P-value < 0.05) test result. This analysis was performed separately for nuclei from the young and aged mice so we could compare the results with respect to aging. We cut the pseudotime trajectory into 10 and 50 evenly sized bins, where each bin contains the same number of cells, for further analysis and data visualization with pseudo-bulk profiles of these bins. We next used a recurrent variational autoencoder (RVAE) ^60^ to compute de-noised gene expression signatures of these t-DEGs in evenly spaced windows throughout the trajectory and to compute a low-dimensional embedding of these genes with respect to their temporal expression patterns. Out of the t-DEGs, we also sought to identify genes with switch-like expression patterns which may be important in the transition phase between progenitors and oligodendrocytes. We defined “switch genes” as genes that reached 80% of its maximum expression level in only two out of 50 trajectory windows.

### Gene co-expression network analysis in the oligodendrocyte lineage

We performed weighted gene co-expression network analysis (WGCNA)^56^ in the OLCs of the snRNA-seq dataset using the R package hdWGCNA^20^ (v 0.4.02). Prior to network analysis, we computed pseudo-bulk gene expression profiles for each of the snRNA-seq samples split by the 10 pseudotime windows using the hdWGCNA function ConstructPseudobulk followed by a log2(CPM) normalization. We used the set of oligodendrocyte trajectory DEGs t-DEGs from both the young and old mice as the input features for network analysis. Using the hdWGCNA function TestSoftPowers, we tested the scale-free topology model fit for different soft-power thresholds and selected the smallest power that yielded a fit greater than 0.8. We then constructed the co-expression network using the hdWGCNA function ConstructNetwork with a minimum module size of 50, a merge cut height of 0.05 and a detect cut height of 0.995 for Dynamic Tree Cutting^57^, and default values for all other parameters. From this analysis we obtained 10 co-expression modules and a topological overlap matrix (TOM) containing co-expression information for each pair of genes. We computed MEs to summarize gene expression information for each co-expression module in each cell using the hdWGCNA function ModuleEigengenes, and we applied Harmony correction to the MEs based on sequencing batch. We then computed eigengene-based connectives (kMEs) for each gene using the hdWGCNA function ModuleConnectivity, which thereby allows us to identify the most connected genes in each module, also known as the module hub genes.

By inspecting the gene expression patterns of each module throughout the trajectory and with respect to the distinct subtypes within the OLCs (OPCs, MFOLs, etc), we noted that one of the largest modules, M2, was exclusively expressed in the OPCs. To identify additional patterns within this module, we performed another round of network analysis using the genes in module M2 in the snRNA-seq nuclei annotated as OPCs. We performed this additional network analysis with hdWGCNA using the same functions and settings as previously described above, identifying 18 additional co-expression modules specific to OPCs. We identified enriched biological processes in each of the co-expression modules using the R package EnrichR (v 3.1) using the following reference databases: GO Biological Processes 2021, GO Cellular Component 2021, GO Molecular Function 2021. We visualized the gene expression patterns of the MEs throughout the oligodendrocyte lineage trajectory with the hdWGCNA function PlotModuleTrajectory, and we plotted heatmaps showing the gene expression trajectories of individual gene faceted by co-expression module. We performed differential module eigengene (DME) analysis with the hdWGCNA function FindDMEs to compare the co-expression module expression patterns between young and aged mice throughout the oligodendrocyte lineage trajectory. We assessed the epigenomic activity of these co-expression modules by projecting the modules into the snATAC-seq dataset. We used the hdWGCNA function ProjectModules to calculate MEs in the OLCs from the snATAC-seq dataset using the ArchR gene score assay.

### Transcription Factor Regulatory Network Analysis of the Oligodendrocyte Lineage

We used the information from our integrated single-nucleus multi-omic oligodendrocyte lineage to perform transcription factor (TF) regulatory network analysis. The goal of this analysis is to identify active trans-regulatory links between TFs and their putative target genes. Here we use a custom bioinformatic pipeline leveraging the integrated gene expression and chromatin accessibility information to infer TF regulatory networks, with the full code and documentation available on our GitHub page. Importantly, we performed the TF regulatory network analysis separately on nuclei from the young and aged populations to facilitate downstream comparisons. Additionally, we selected specific cell populations in our dataset for this analysis to capture the regulatory changes underlying re-myelination: Since TFs can regulate target genes through binding sites at a gene’s promoter or at a distal enhancer binding site mediated by chromatin looping, we first collected information about genome-wide motif binding sites using the R package motifmatchr (v 1.12.0) to search for position weight matrices (PWMs) for TF binding motifs in the JASPAR2020^48^ database. We established links between TFs and putative target genes if there was a TF motif present in a gene’s promoter region or if a TF was present in a linked distal cCRE from our previously described chromatin co-accessibility analysis, thereby capturing enhancer mediated TF-gene regulatory interactions. We used these initial TF-gene mappings to further identify highly confident regulatory relationships by modeling each gene’s expression based on the expression of its potential TF regulators. We next used extreme gradient boosting trees (XGBoost) to build an ensemble of regression models to predict the gene expression of each gene based on the expression levels of their putative TF regulators. For this analysis we performed 5-fold cross validation and averaged the performance statistics across these folds. XGBoost regression provides us with information about how important each TF was for accurately predicting the expression of a particular gene, therefore allowing us to systematically prioritize which TFs were likely actively regulating each gene. Using this information we defined sets of confident target genes for each TF, called TF regulons, by retaining the top five TFs for each gene based on their XGBoost importance scores with a minimum threshold of 0.001. We further stratified these TF regulons based on gene expression correlations between the TF and the target genes, where positive correlations represent putative “activating” relationships and negative correlations represent putative “repressing” relationships. Using the TF motif binding information and the results from our XGBoost regression analysis, we construct TF regulatory networks where nodes in the network represent genes or TFs and directed edges represent with a weighted regulatory importance score. We used the R package igraph (v 2.0.3) to assemble these networks and for downstream data visualization. Together with the enhancer and promoter binding site information, TF regulatory network analysis provides potential mechanistic understanding for gene regulatory relationships that can be further probed with experimental validation.

### Optimal pathfinding to identify cascades of transcription factor regulation

We sought to use the TF regulatory networks from young and aged mice along with the multi-omic oligodendrocyte trajectory to computationally infer the temporal dynamics of TF binding and subsequent gene expression changes in remyelination, and to understand how these dynamics change between young and aged mice. For this analysis, we focus only on the TF-TF regulatory relationships in our networks. In essence, our goal is to identify a core directed subgraph within these TF-TF networks that represent the most critical regulators of remyelination and oligodendrocyte maturation. Several key TFs are already known to be involved in these processes based on previous studies, including *Sox10*, *Nkx2-2*, *Nkx6-2*, *Tcf4*, *Olig1*, and *Olig2*, and we leverage this prior knowledge in our approach.

Here we employed Dijkstra’s algorithm^58^ to perform optimal pathfinding analysis from TFs highly expressed in OPCs at the earliest point in the pseudotime trajectory (start nodes) towards those most highly expressed in Mat-ODCs at the terminus of the pseudotime trajectory (end nodes) to predict potential TF regulatory cascades. For this pathfinding analysis, we used the subgraph of the full TF regulatory network containing only TFs and their links with other TFs, excluding all other genes. We weighted the edges of this network with a custom score accounting for binding site information (TF motif present at target’s promoter, enhancer, or both), the regulatory importance score from our XGBoost regression analysis, TF-TF co-expression, and the pseudotime bin distance. Since Dijkstra’s algorithm identifies optimal paths in a network by traversing low-cost edges, small but positive edge weights in our network represent TF-TF regulatory relationships with strong evidence from multiple sources. Importantly, this analysis does not account for repressive regulatory relationships, since only edges with positive weights are retained for analysis. We first identified the shortest paths in this network between the starting nodes and the six previously mentioned TFs with substantial prior evidence for their involvement in oligodendrocyte differentiation, which we termed “critical nodes” in this network. We next identified the shortest paths between the critical nodes and the end nodes. To consider TF cascades that do not involve the critical nodes, we also identified the shortest paths between the start nodes and the end nodes. We finally assemble the nodes and edges of these individual paths into a new graph. We visualized this directed graph in the RVAGene latent space to visually distinguish the pseudotemporal relationships between TFs throughout oligodendrocyte maturation. We computed key network statistics like centrality and degree within this subgraph. We implemented this pathfinding analysis using the R package igraph.

### Reprocessing published human MS snRNA-seq datasets

We collected raw sequencing data (.fastq format) for two previously published snRNA-seq datasets from human MS donors and cognitively normal controls. The Absinta et al 2021^33^ snRNA-seq dataset was collected from the Gene Expression Omnibus (GEO) at accession GEO180759, and the Schirmer and Velmeshev et al 2019^32,33^ dataset was collected from the Sequence Read Archive (SRA) at accession PRJNA544731. For each biological replicate, we used Kallisto Bustools^59^ to pseudoalign raw sequencing reads to the human reference transcriptome (GRCh38) and quantify gene expression levels for each cell barcode. The human reference transcriptome was obtained from the 10X Genomics website (v 2020-A, July 2020), and was re-formatted for use with Kallisto Bustools using the kb ref function. We then ran CellBender^60^ to correct ambient RNA contamination in each of the snRNA-seq samples. Similar to our mouse snRNA-seq dataset, we ran Scrublet to identify and remove potential doublets from these datasets. Individual counts matrices were then merged into a single anndata object for the two datasets for downstream analysis with the Python package Scanpy (v 1.6)^61^.

We performed the subsequent clustering analyses separately for the Absinta et al. 2021 and the Schirmer and Velmeshev et al. 2019 datasets using a similar processing pipeline; the exact code and parameters used for this analysis can be viewed on our GitHub. Prior to clustering, we first performed a percentile filtering of outlier cells in each sample based on the number of UMI per cell, the percentage of UMI attributed to mitochondrial genes, and the predicted doublet score from Scrublet. After filtering low quality cells, we calculated log-normalized gene expression profiles using the Scanpy functions sc.pp.normalize_total and sc.pp.log1p. We selected highly variable genes for downstream dimensionality and clustering analysis using the Scanpy function sc.pp.highly_variable_genes. The normalized gene expression matrix was then scaled to unit variance and centered at zero using the Scanpy function sc.pp.scale. Principal component analysis (PCA) was then used to linearly transform and reduce the dimensionality of the scaled expression matrix while retaining information content, and Harmony was used to correct the resulting PCA representation for batch effects with the Python package harmonypy. We then constructed a cell neighborhood graph based on the harmonized PCA matrix using the Scanpy function sc.pp.neighbors, which was then used for Leiden clustering (sc.tl.leiden function) and UMAP (sc.tl.umap). We assigned coarse-grain cell type labels to each of the Leiden clusters by inspecting gene expression signatures for a panel of canonical cell type marker genes in each cluster. At this stage, we also inspected the distribution of QC metrics in each cluster in order to identify clusters of low-quality cells or outliers for removal from subsequent analyses. We filtered out additional outlier clusters based on these QC metrics or if the cluster had conflicting marker gene expression, indicating a doublet cluster that was not previously identified by Scrublet. After this additional filtering step, we re-computed the Leiden clusters and the UMAP, resulting in a final processed dataset. We then used a custom script to convert the dataset from anndata to SeuratObject format for downstream analysis.

### Differential gene expression analysis in the human MS datasets

We performed a series of differential gene expression analyses in the re-processed Absinta et al. 2021 and Schirmer and Velmeshev et al. 2019 snRNA-seq datasets. Importantly, these studies profiled gene expression in single nuclei from MS patients in several different sites with respect to MS lesions. The Absinta et al. 2021 dataset included snRNA-seq data for chronic active MS lesion edge, chronic inactive MS lesion edge, MS lesion core, MS periplaque WM, and WM from cognitively normal control donors. The Schirmer and Velmeshev et al. 2019 dataset used snRNA-seq to profile cortical grey matter (GM) and adjacent WM MS lesions, with data for acute chronic active MS lesions, chronic inactive MS lesions, and control tissue. Similar to our previously described differential expression analysis with the mouse dataset, we used MAST as the underlying statistical model, and we used the following variables as model covariates: sample ID, number of UMI per cell, mitochondrial count percentage, postmortem interval (PMI), and RNA integrity number (RIN). For the Absinta et al. 2021 dataset we performed the following five differential expression tests: chronic active MS lesion edge versus control WM, chronic inactive MS lesion edge versus control WM, chronic active MS lesion edge versus chronic inactive MS lesion edge, MS lesion core versus control WM, and MS periplaque WM versus control WM. For the Schirmer and Velmeshev et al. 2019 dataset we performed the following three differential expression tests: acute chronic active MS lesion versus control, chronic inactive MS lesion versus control, and acute chronic active MS lesion versus chronic inactive MS lesion.

### Data driven prioritization of TFs for experimental validation

We used a holistic data-driven approach to nominate TFs of interest for additional experimental validation. This approach leveraged information from multiple of our previously described data analysis steps. We set the following criteria for potential TFs of interest: 1, DEGs in the mouse dataset that are differentially expressed between young and aged mice in oligodendrocytes; 2, DEGs in the human dataset to identify genes that are differentially expressed between MS and control patients; 3, genes intersecting between these mouse and human DEGs; 4, t-DEGs in young or aged mice; 5, t-DEGs with switch-like gene expression patterns (not used as a strict filter); 6, genes in the core TF regulatory network from our pathfinding analysis. Based on these criteria we selected seven TFs for downstream experiments: *Nr6a1* (GCNF), SOX8, FOXK2, BHLHE41, STAT3, BACH2, and ELF2.

### Isolation of primary rodent OPCs

Neonatal OPCs (P7-P10) were isolated as previously described^44,62^. In brief, rat pups were euthanized with an overdose of sodium pentobarbital. Brains were dissected, the meninges removed and the brains were finely chopped, before enzymatic digestion with papain and DNase type I. After digestion, the tissue suspension was further dissociated by tritruation with fire-polished Pasteur pipettes. The Homogeneate was filtered through a 70 µm cell strainer and centrifugated in isotonic 22.5% Percoll (800 g, 20 min, RT, no break). The resulting cell pellet was briefly washed in Hanks’ Balanced Salt Solution (HBSS) and resuspended in red blood cell lysis buffer (Sigma, R7757) for 1 minute at RT. After red blood cell lysis 2.5 μg of a magnetic bead-conjugated mouse-anti-rat-CD45 antibody (Miltenyi) was added with 2.5 μg of mouse anti-A2B5 antibody to the cell suspension for simultaneous negative selection of CD45+ cells and primary labelling of OPCs with A2B5 antibody. Magnetic-activated cell sorting (MACS, Miltenyi) following the recommendations of the manufacturer was performed for negative selection.The flow-through containing A2B5-labelled cells was briefly washed and subsequently incubated with magnetic bead-conjugated rat-anti-mouse-IgM antibody (Miltenyi). Following incubation, another round of MACS was performed according to the manufacturers instruction, and CD45-/A2B5+ cells were eluted with pre-equilibrated OPC media. Cells were seeded onto poly-D-lysine coated cell-culture treated plates and maintained in complete OPC media (60 μg/mL N-acetyl cysteine (Sigma), 10 μg/mL human recombinant insulin (GIBCO), 1 mM sodium pyruvate (GIBCO), 50 μg/mL apo-transferring (Sigma), 16.1 μg/mL putrescine (Sigma), 40 ng/mL sodium selenite (Sigma), 60 ng/mL progesterone (Sigma), 330 μg/mL bovine serum albumin (Sigma)), supplemented with basic fibroblast growth factor (b-FGF) and platelet-derived growth factor (PDGF, 30 ng/mL each, Peprotech)). For differentiation conditions, cells were transferred into media lacking the mitogens PDGF and FGF. Differentiation enhancing 40 ng/mL thyroid-hormone (T3, Sigma) was added for positive controls in the differentiation assays. Cultures were maintained throughout at 37°C, 5% CO_2_ and 5% O_2_.

### Cell culture and lentiviral infection of OPCs

Lentiviral particles containing the transgene of interest under the mouse PGK promoter together with a CMV-driven GFP:T2A:puromycin selection cassette (Vectorbuilder) were added to the OPCs in complete OPC media for two days, followed by a full media change with complete OPC media with b-FGF and PDGF. The cells were incubated for another 2 days to allow expression of transgenes and successfully infected, GFP expressing cells were subsequently selected with 1 μg/mL puromycin (ThermoFisher A1113803) in complete OPC media containing b-FGF and PDGF for 2 days. After 2 days virtually all remaining cells were GFP positive and either processed for proliferation or differentiation assays. For proliferation assays, OPCs were fixed with 4% PFA in PBS for 15 minutes, washed 3 times with PBS and stored at 4°C for immunocytochemistry for Ki-67. For differentiation, OPCs were changed into OPC media without miogens. T3-treated OPCs served as a positive control. Cells were allowed to differentiate for 3 days for immunocytochemistry, after which they were fixed in 4% PFA in PBS, washed 3 times with PBS and stored at 4°C until processed for immunocytochemistry. For bulk RNAsequencing, cells in differentiation condition underwent mitogen withdrawal for 24h.

### Immunocytochemistry, microscopy, and quantification

To assess OPCs for proliferation and differentiation, cells were first washed twice with PBS and then blocked with a blocking buffer (10% normal donkey serum in PBS with 0.1% Triton X-100) for 1 hour at room temperature. Cells were then incubated with primary antibodies for 2 hours at room temperature. To identify proliferating OPCs, a rat anti-Ki67 (Thermofisher 14-5698-82, clone SolA15) antibody and a rabbit anti-Olig2 (Merck, ab9610) antibody were used. To investigate the extent of OPCs differentiating into oligodendrocytes, a rat anti-MBP (Serotec/Biorad) antibody and a rabbit anti-Olig2 (Merck, ab9610) antibody were used. After incubation with the primary antibodies, cells were washed 3 times with PBS, followed by incubation with appropriate secondary antibodies for 2 hours at room temperature. Cells were then washed 3 times with PBS and then stored at 4°C in the dark in PBS.

Plates were imaged on a high-throughput Opera imaging system, taking 69 fields of view per well to capture the entire well with z-stacks. Maximum intensity projections were produced for quantification. All images were blinded for quantification. CellProfiler was used to quantify the percentage of Olig2+ cells that were Ki67+ (proliferation) and the percentage of Olig2+ cells that were MBP+ (differentiation). To assess the effect of transcription factor overexpression on proliferation and differentiation rates, linear mixed-effects models were fitted using the lme4 package in R (4.4.2). The response variables were the logit-transformed percentages of proliferating and differentiating cells (3 technical replicates, 3-4 biological replicates per condition), or the absolute counts of OLCs (Olig2+, n=3 biological replicates per condition), as indicated. Transcription factor overexpression was included as a fixed effect, and a random intercept was added to account for inter-animal variability. P-values for the fixed effect were generated via Satterthwaite’s method, adjusted for multiple comparisons using the Benjamini-Hochberg method, and significance was determined at an FDR-adjusted q < 0.05.

### Bulk RNA sequencing processing

Cells in proliferation or differentiation conditions were washed twice with PBS and collected in Trizol. Library preparation was performed using the NEB NEBNext Ultra II Directional RNA kit (New England Biolabs) on a Firefly instrument (SPT Labtech Ltd.). Library preparation was performed following the manufacturer’s recommendations and sufficient RNA quality was confirmed by Tapestation measurement. The sequencing was performed on an Illumina Novaseq platform.

RNA-seq data was processed using the nf-core/rnaseq pipeline^63^ (v3.12.0) implemented in Nextflow^64^ (version 23.10.1). Raw sequencing reads were subjected to quality trimming and adapter removal using Trim Galore (v0.6.7) with Cutadapt^65^ (v3.4). Read alignment against the rat mRatBN7.2 genome was performed using STAR^66^ (v2.7.9a) with default parameters optimized for RNA-seq data using transcript models based on Ensembl release v108.

Gene and transcript quantification was performed using Salmon^67^ (v1.10.1) in quasi-mapping mode. For alignment-based quantification, feature counting was performed using Subread’s featureCounts^68^ (v2.0.1). Comprehensive quality control reports were generated using MultiQC^69^. Reads aligned to the genome by STAR were further visualized and quantified using SeqMonk (v1.48.0) (https://github.com/s-andrews/SeqMonk). . Statistical analysis of differential gene expression was carried out in R(4.4.0) using the limma and voom packages. Genes were filtered using the filterByExpr function in edgeR with default settings (min.count = 10, min.total.count=15). A linear model was fit for each gene using both transcription factor overexpression and animal as factors. Obtained p-values were adjusted for multiple comparisons using the Benjamini-Hochberg method, and genes were determined as differentially expressed at an FDR-adjusted q < 0.05.

### Data availability

All raw and processed single-nucleus sequencing data have been deposited into the National Center for Biotechnology Information Gene Expression Omnibus under accession number GSE277068. At this time, data access remains private and the peer reviewers have been provided with a secure access token. Additional snRNA-seq datasets from published studies were obtained via the following accessions: Absinta et al 2021^33^ snRNA-seq dataset was collected from the Gene Expression Omnibus at accession GEO180759, and the Schirmer and Velmeshev et al 2019^32,33^ dataset was collected from the Sequence Read Archive (SRA) at accession PRJNA544731.

### Code availability

The data analysis code used for this study is available on GitHub at https://github.com/smorabit/Myelination_2025.

## Acknowledgements

R.J.M.F and V.S. acknowledge the Dr Miriam and Sheldon G Adelson Medical Research Foundation for support. R.J.M.F was supported by a core support grant from the Wellcome Trust and MRC to the Wellcome Trust-Medical Research Council Cambridge Stem Cell Institute (203151/Z/16/Z) and funding from the UK Multiple Sclerosis Society (MS50). V.S. was supported by National Institute on Aging grants 1R01 AG071683, 1R01 NS135556, 2U54AG054349 (MODEL-AD), and 1R01AG082147-01; National Institute of Neurological Disorders and Stroke grant 1R01 NS135556. S.M was supported by a National Institute on Aging predoctoral fellowship F31AG076308-01 and a European Molecular Biology Organization postdoctoral fellowship ALTF-834-2024. K.S.R was supported by a Banting Postdoctoral Fellowship from the Canadian Institute of Health Research as well as a Multiple Sclerosis Society of Canada Fellowship. P.A. was funded by a Banting Postdoctoral Fellowship from the Canadian Institute of Health Research. We acknowledge funding from the European Union’s H2020 research and innovation program (grant agreement 848028) and from the European Research Council (ERC) under the European Union’s Horizon 2020 Research and Innovation Program (grant agreement 810287) awarded to H.H. Further support to H.H. was received from the Ministerio de Ciencia e Innovación (MCI) under grant agreements PID2020-115439GB-I00 and PLEC2021-007654, and by ERA-NET Neuron/Ministerio de Ciencia e Innovación (MCI) under grant agreement PCI2022-133012. H.H. has received additional funding from the European Union’s Horizon Europe Research and Innovation program (grant agreement 101072891) and from LaCaixa Foundation (grants HR22-0031 and HR22-0172). We are also thankful to the Generalitat de Catalunya through the Suport Grups de Recerca AGAUR (2021-SGR2021-01216 to H.H.). This work used the infrastructure for high-performance and high-throughput computing, research data storage and analysis, and scientific software tool integration built, operated and updated by the Research Cyberinfrastructure Center at the University of California Irvine. We acknowledge the Altos Hub facility contributions of Dr. Felix Krueger, Dr. Christel Krueger, Dr. Paul Coupland, and Dr. Hanneke Okkenhaug. We are also grateful to Dr. Jonathan Griffiths for his statistical insight and advice.

## Author contributions

P.D., S.M., K.S.R., R.J.M.F, and V.S. wrote the manuscript with assistance from all authors. P.D., K.S.R., and J.C. performed the mousework and Q.W. and R.K. generated the single-nucleus sequencing data. S.M. conducted bioinformatics analysis with assistance from Z.S. and Z.C. P.D. and K.S.R. performed the *in vitro* experiments with help from P.A. and N.B. F.N.K. conducted the statistical analysis of the *in vitro* experiments.

