## Supplementary Materials for "Dysregulation of transcription networks regulating oligodendrogenesis in age-related decline in CNS remyelination"

### Supplementary Tables

**Supplementary Table 1**: Samples used for snRNA-seq and snATAC-seq, and summary of clusters
**Supplementary Table 2**: snRNA-seq cluster marker genes
**Supplementary Table 3**: Oligodendrocyte trajectory differentially expressed genes (t-DEGs) in young mice.
**Supplementary Table 4**: Oligodendrocyte trajectory differentially expressed genes (t-DEGs) in aged mice.
**Supplementary Table 5**: Oligodendrocyte co-expression network modules from hdWGCNA.
**Supplementary Table 6**: Oligodendrocyte progenitor co-expression network modules from hdWGCNA.
**Supplementary Table 7**: Inferred transcription factor (TF) target genes (regulons) from TF regulatory network analysis in the oligodendrocyte lineage in young mice.
**Supplementary Table 8**: Inferred transcription factor (TF) target genes (regulons) from TF regulatory network analysis in the oligodendrocyte lineage in aged mice.
**Supplementary Table 9**: Differential gene expression results comparing nuclei from Aged vs. Young in each snRNA-seq cluster.
**Supplementary Table 10**: Differential gene expression results comparing nuclei across disease-relevant samples in each snRNA-seq cell type in the human datasets.
**Supplementary Table 11**: Bulk RNA-seq differential expression results.

### Supplementary Figures

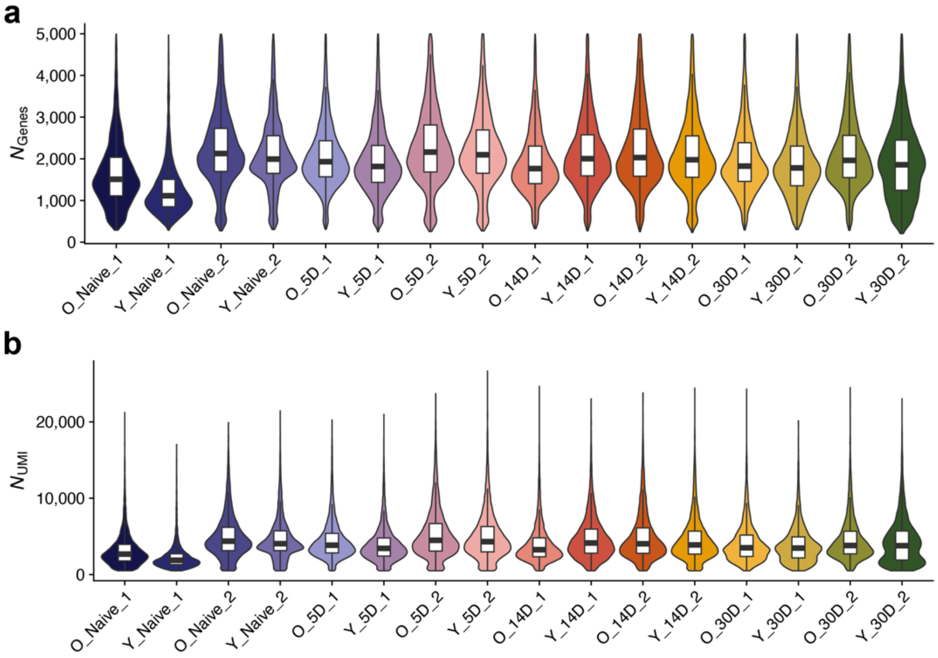


**Supplementary Fig. 1 | Quality control metrics in the snRNA-seq dataset.** Violin plots showing the distribution of quality control (QC) metrics in each sample used for snRNA-seq. **a**, The number of genes detected in each nucleus. **b**, the number of unique molecular identifiers (UMI) detected in each nucleus. For boxplots, box boundaries and lines correspond to the interquartile range (IQR) and median respectively. Whiskers extend to the lowest or highest data points that are no further than 1.5 times the IQR from the box boundaries

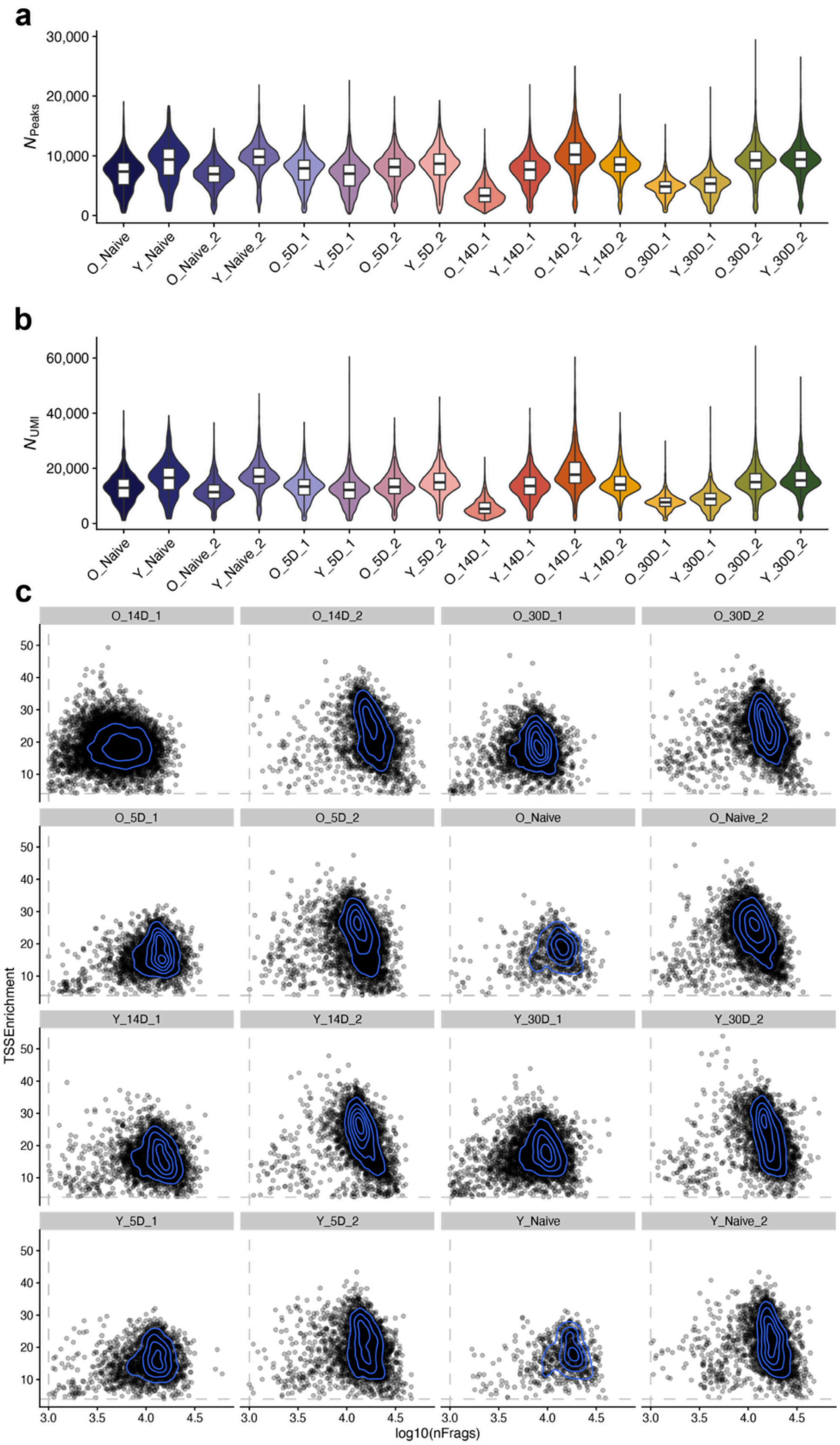


**Supplementary Fig. 2 | Quality control metrics in the snATAC-seq dataset.** Violin plots showing the distribution of quality control (QC) metrics in each sample used for snATAC-seq. **a**, The number of ATAC peaks (features) detected in each nucleus. **b**, the number of unique molecular identifiers (UMI) detected in each nucleus. For boxplots, box boundaries and lines correspond to the interquartile range (IQR) and median respectively. Whiskers extend to the lowest or highest data points that are no further than 1.5 times the IQR from the box boundaries. **c**, Density scatter plots comparing the number of unique ATAC fragments vs. transcription start site (TSS) enrichment scores in each snATAC-seq nucleus, faceted by each sample. The QC threshold used for the downstream analysis are indicated by the grey dotted lines.

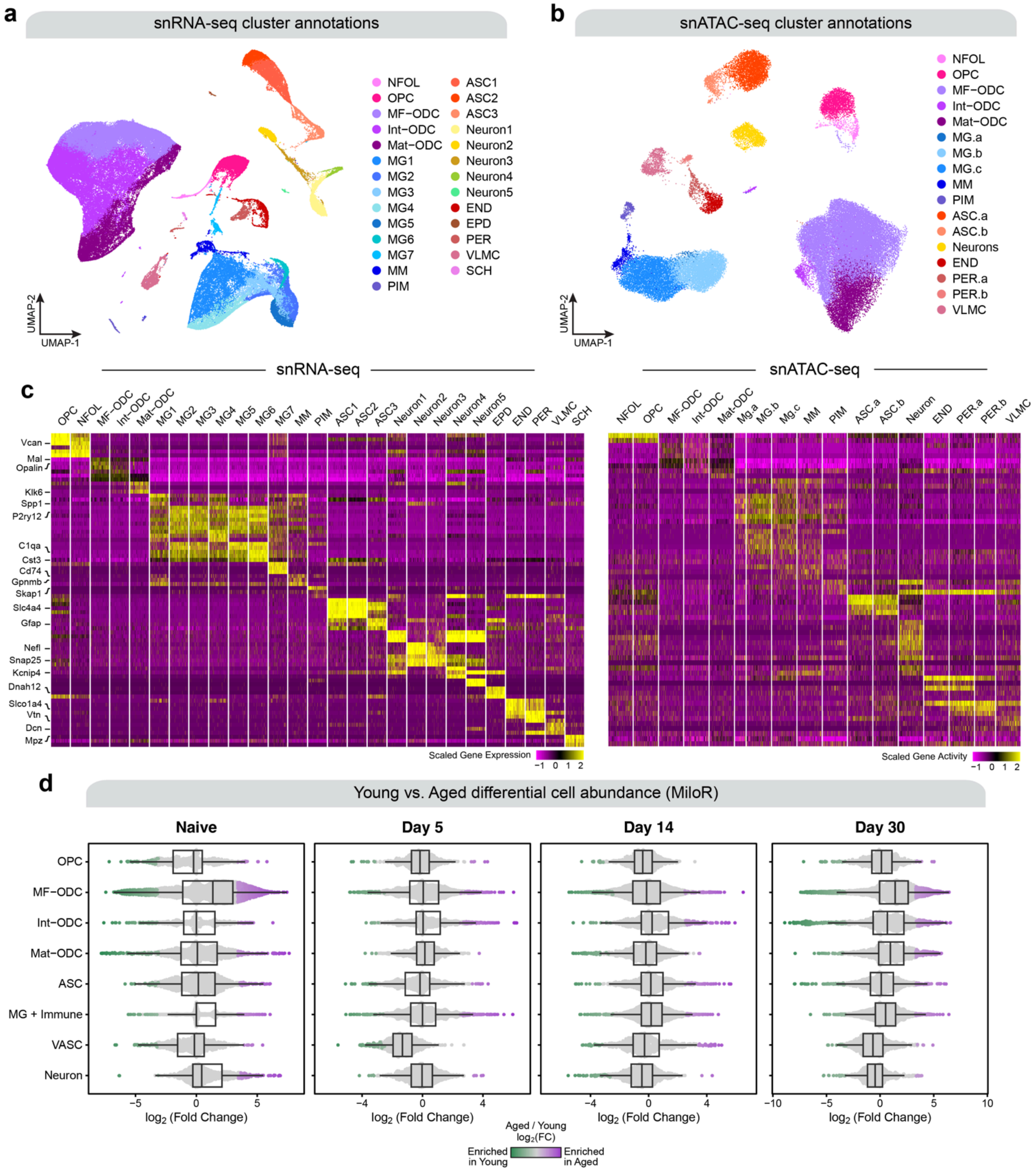


**Supplementary Fig. 3 | Individual clustering analyses of the snRNA-seq and snATAC-seq datasets. a**, UMAP plot of 157,563 snRNA-seq profiles. The UMAP layout is based on Liger[1] integrative non-negative matrix factorization (iNMF) dimensionality reduction of snRNA-seq gene expression. Cells are colored by annotated snRNA-seq clusters. **b.** UMAP plot of 52,688 snATAC-seq profiles. The UMAP layout is based on iterative latent semantic indexing (iLSI) dimensionality reduction with Harmony[2] correction of the snATAC-seq chromatin accessibility tiles. Cells are colored by annotated snATAC-seq clusters. **c**, Left: heatmap showing the scaled gene expression of the top three marker genes by average $\log_2$(fold change) from each of the 29 snRNA-seq clusters. Right: heatmap of the same genes shown in the scaled gene activity matrix from the snATAC-seq dataset. **d.** Differential cell abundance results comparing aged vs. young within the four experimental timepoints. Box boundaries and lines correspond to the IQR and median, respectively. Whiskers extend to the lowest or highest data points that are no further than 1.5 times the IQR from the box boundaries. Each data point represents a single-cell neighborhood from Milo[3]. Neighborhoods with a significant difference ($P$-value < 0.05) are colored based on the $\log_2$(fold change).

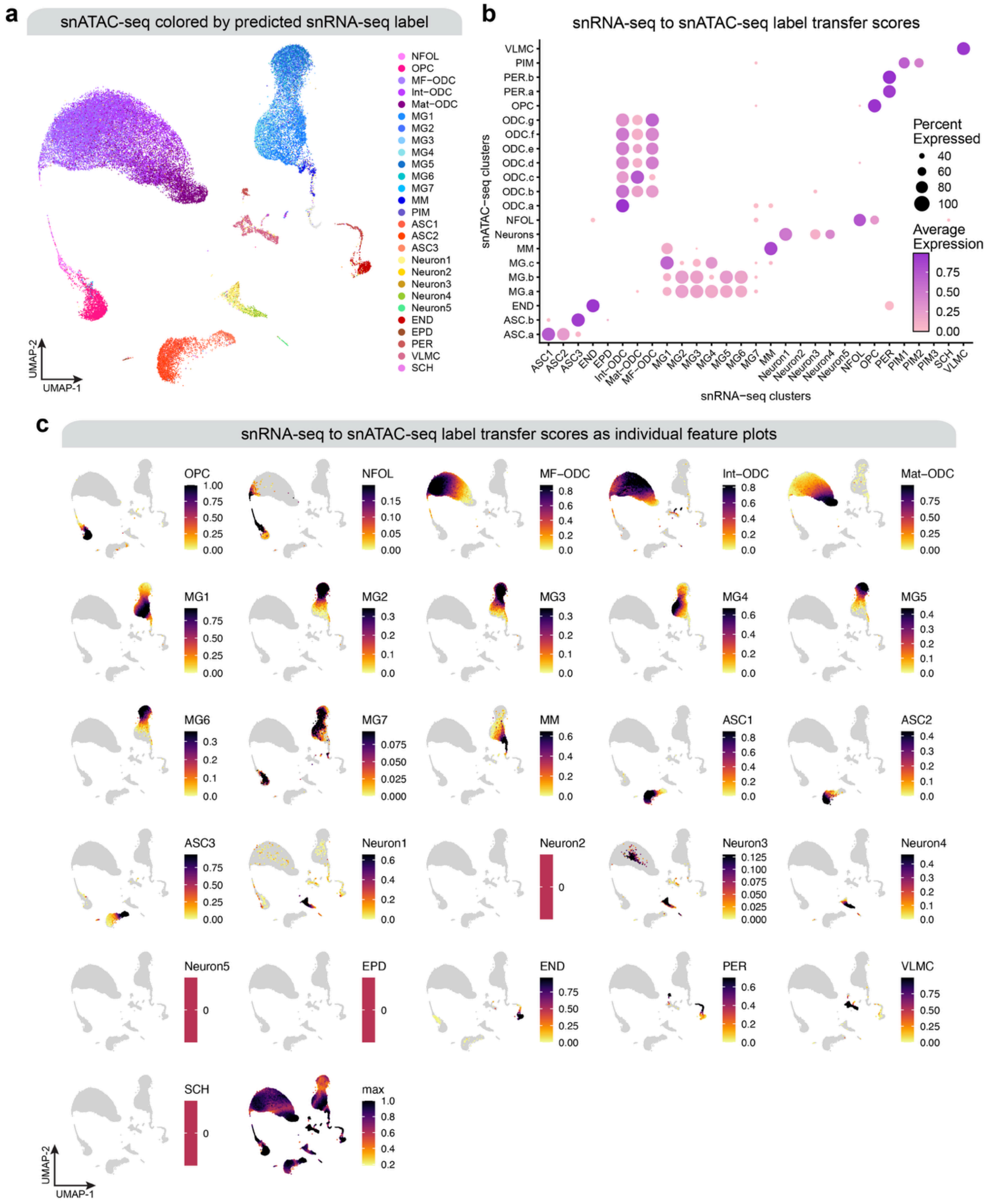


**Supplementary Fig. 4 | Integration and label transfer between snRNA-seq and snATAC-seq. a,** UMAP plot of 52,688 snATAC-seq cells. The UMAP was calculated from the integrated snRNA-seq + snATAC-seq dataset, but only the snATAC-seq data is shown here. snATAC-seq cells are colored by the most likely predicted cell type based on the label transfer analysis with the snRNA-seq dataset. **b**, Dot plot showing the continuous label transfer scores for each of the snRNA-seq clusters in the snATAC-seq. Continuous label transfer scores are calculated in each snATAC-seq cell for each of the 29 snRNA-seq clusters. **c**, UMAP feature plots for the snATAC-seq dataset showing the same label transfer scores as in panel (**b**).

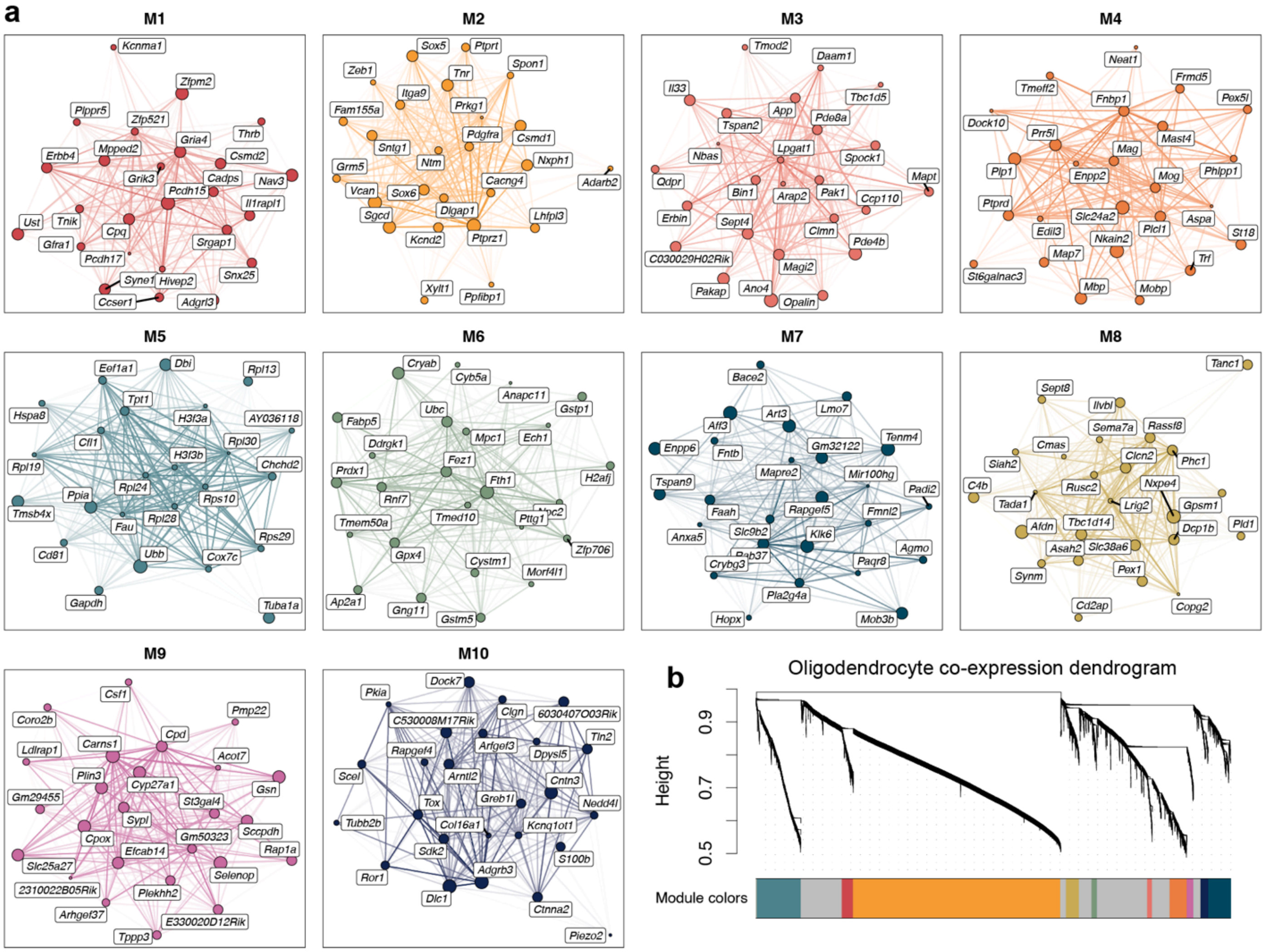


**Supplementary Fig. 5 | Oligodendrocyte lineage gene co-expression network modules. a**, Hub gene networks for each of the 10 oligodendrocyte lineage co-expression modules from hdWGCNA[4]. Nodes represent genes, and edges represent co-expression links. The top 25 hub genes ranked by intramodular connectivity (kME) are shown, and the 250 strongest co-expression links are shown. **b**, Dendrogram plot showing the hierarchical clustering of genes into 10 co-expression modules based on the underlying co-expression network represented as a gene-gene topological overlap matrix (TOM). Module colors are shown below the dendrogram.

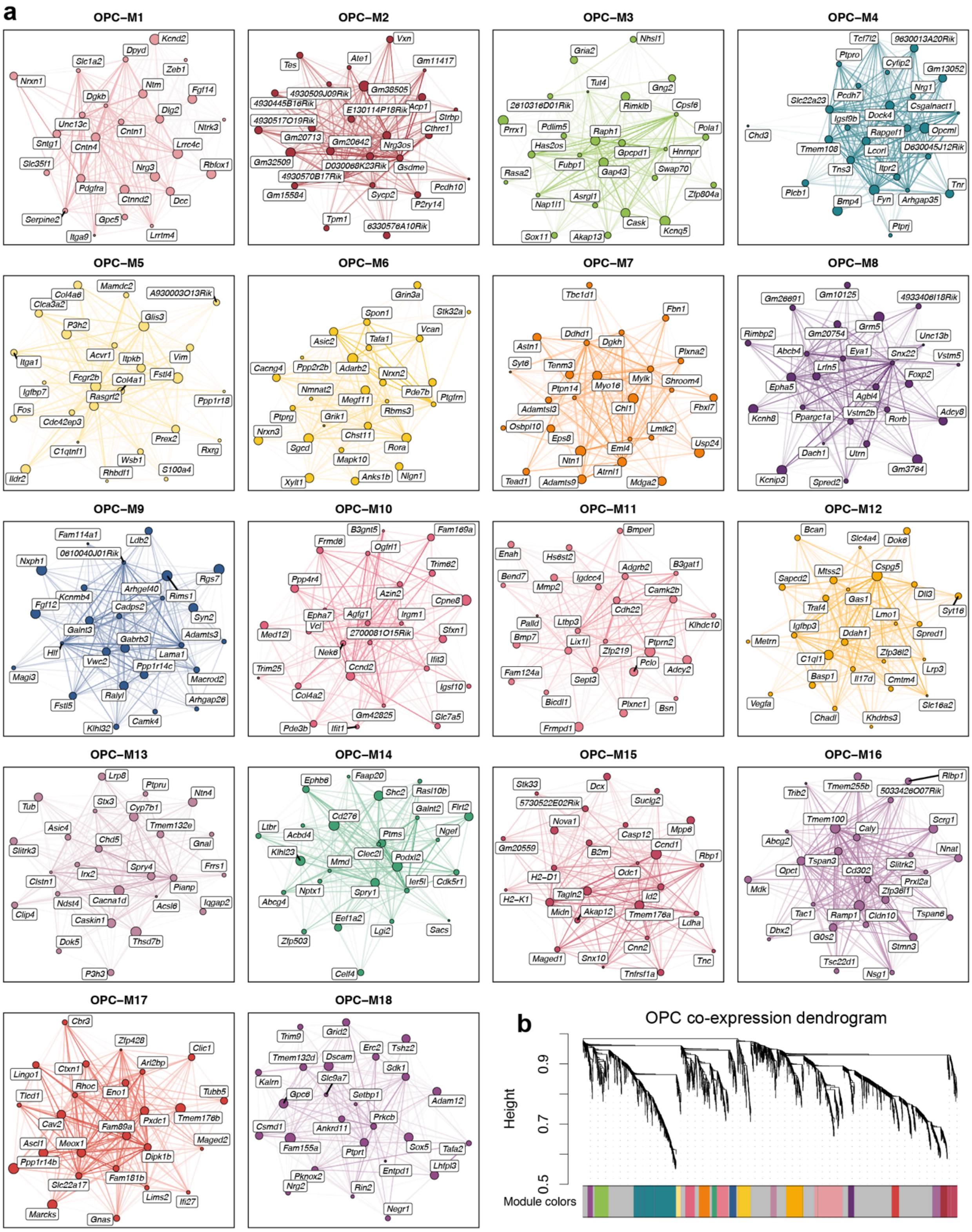


**Supplementary Fig. 6 | Oligodendrocyte progenitor (OPC) gene co-expression network modules. a**, Hub gene networks for each of the 18 oligodendrocyte progenitor co-expression modules from hdWGCNA[4]. Nodes represent genes, and edges represent co-expression links. The top 25 hub genes ranked by intramodular connectivity (kME) are shown, and the 250 strongest co-expression links are shown. **b**, Dendrogram plot showing the hierarchical clustering of genes into 18 co-expression modules based on the underlying co-expression network represented as a gene-gene topological overlap matrix (TOM). Module colors are shown below the dendrogram.

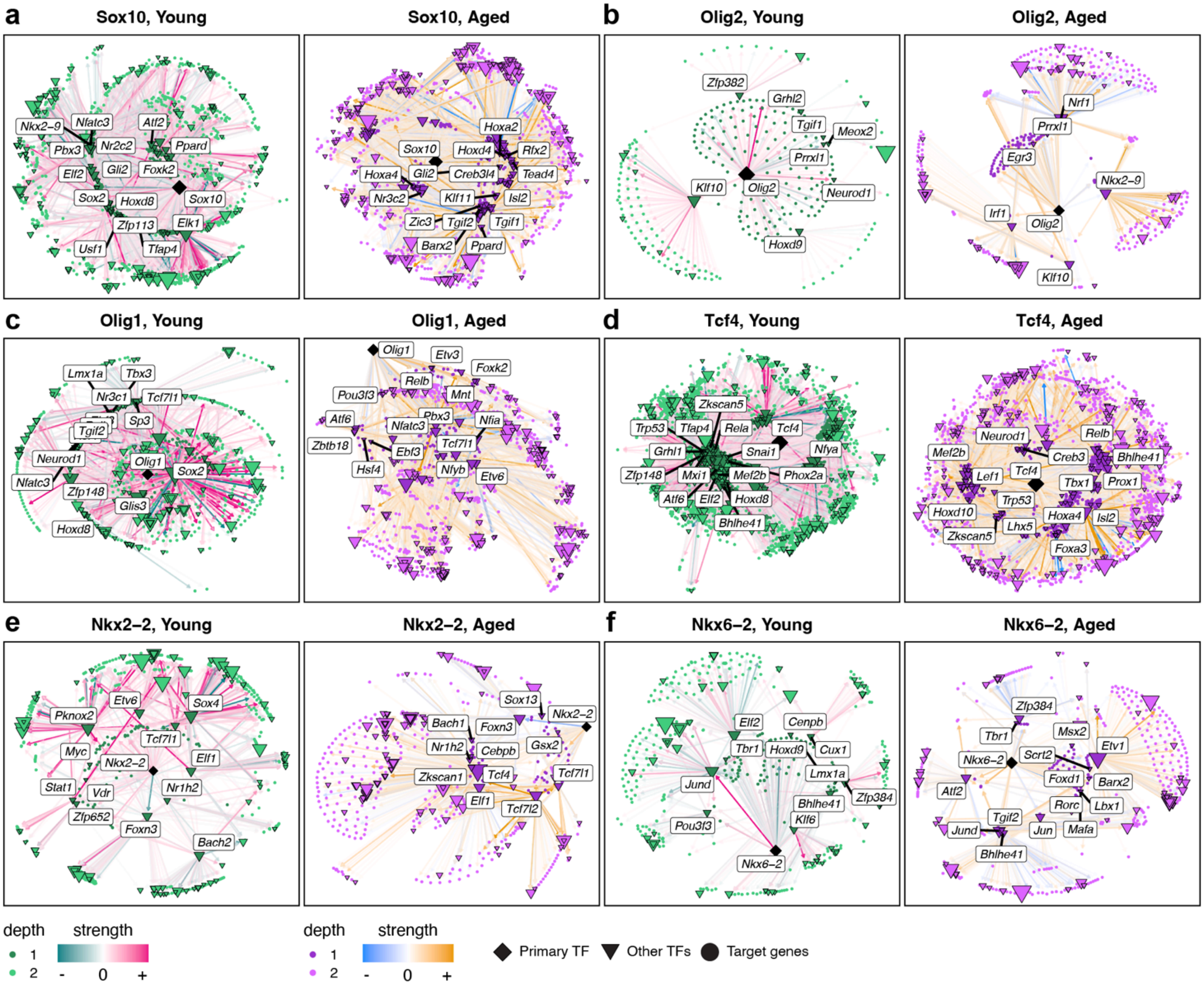


**Supplementary Fig. 7 | Transcription factor regulatory networks in critical oligodendrogenesis TFs.** Transcription factor (TF) regulatory network in young and aged mice for the six "critical" TFs highlighted in the optimal pathfinding analysis: *Sox10* (**a**), *Olig2* (**b**), *Olig1* (**c**), *Tcf4* (**d**), *Nkx2-2* (**e**), and *Nkx6-2* (**f**). Nodes represent genes, and edges represent TF-gene regulatory relationships. Primary (direct) target genes representing the regulon of the main TF of interest (for example, *Sox10* in panel **a**) are shown as darker nodes, and secondary target genes (i.e., the regulons of other TFs targeted by the main TF) are shown in a lighter color. Other TFs targeted by the main TF are labeled.

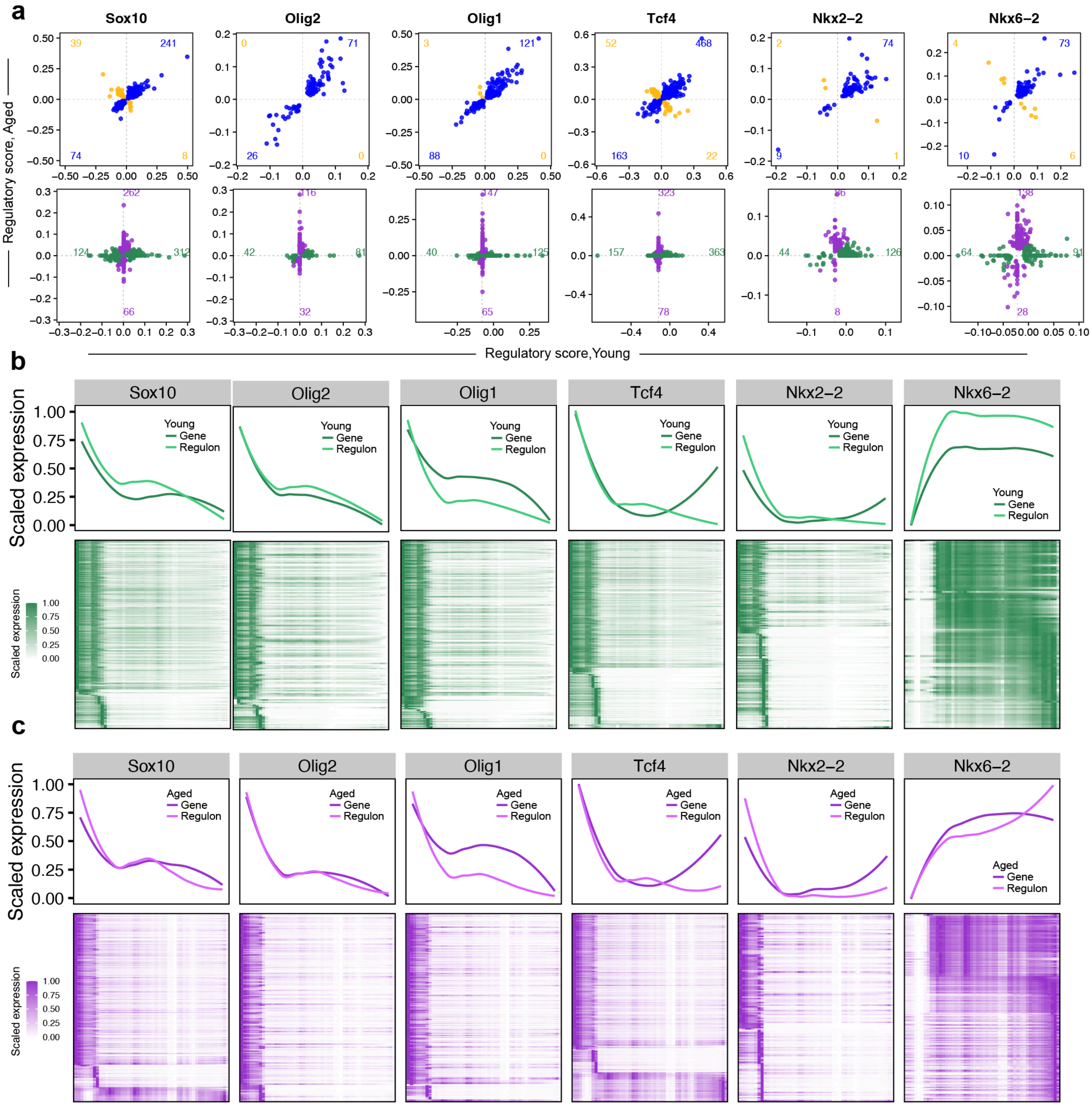


**Supplementary Fig. 8 | Transcription factor expression dynamics in critical oligodendrogenesis TFs. a**, Comparison of TF regulatory scores for the inferred regulons (target genes) of *Sox10, Olig2, Olig1, Tcf4, Nkx2-2,* and *Nkx6-2* in young vs. aged mice. Plots are faceted based on shared genes identified in the regulon in both young and aged (top) and unique genes (bottom). **b**, TF dynamics along the oligodendrocyte pseudotime trajectory in young mice. Top: scaled expression and aggregated TF regulon UCell[5] signature scores along the pseudotime trajectory for the same TFs shown in panel (**a**) Bottom: Heatmaps of scaled gene expression values for the inferred target genes of each TF in young mice. **c**, TF dynamics plots as shown in panel (**b**) for aged mice.

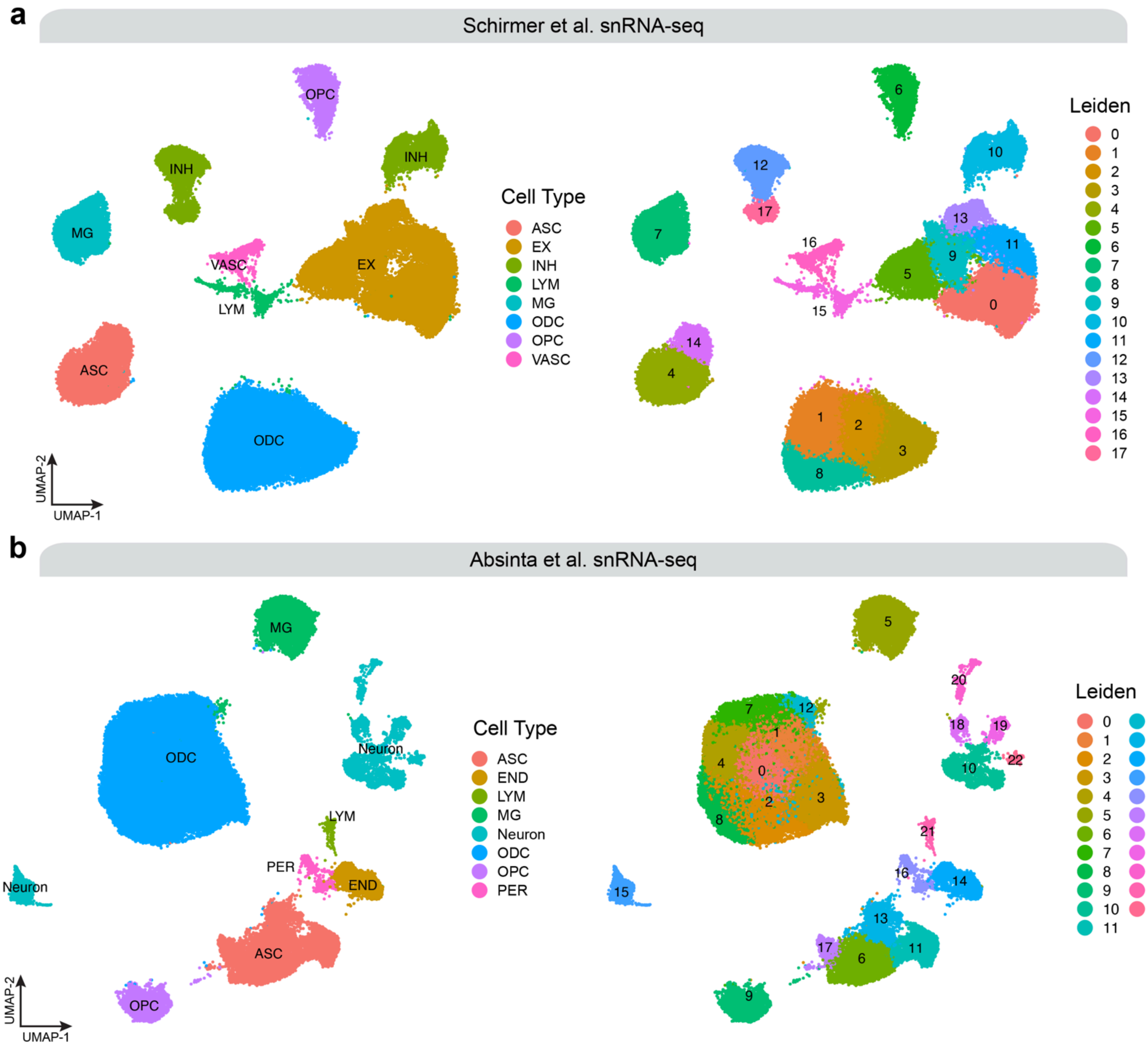


**Supplementary Fig. 9 | Clustering analysis of published human multiple sclerosis snRNA-seq datasets. a**,UMAP plots of 86,578 snRNA-seq profiles from the Schirmer et al 2019[6] dataset. The dataset consists of 8 acute-chronic active multiple sclerosis (MS) lesion samples, 4 chronic inactive MS lesion samples, and 9 control white matter samples. **b.** UMAP plots of 96,865 snRNA-seq profiles from the Absinta et al 2021[7] dataset. The dataset consists of the following samples: 6 chronic active MS lesion edge, 5 chronic inactive MS lesion edge, 2 MS lesion core, 4 MS peri-plaque white matter, and 3 control white matter. Left: cells colored by major cell type annotations. Right: cells colored by unbiased leiden[8] clustering analysis. The raw sequencing data from these two published datasets were re-processed for the purpose of the current study (see Methods).

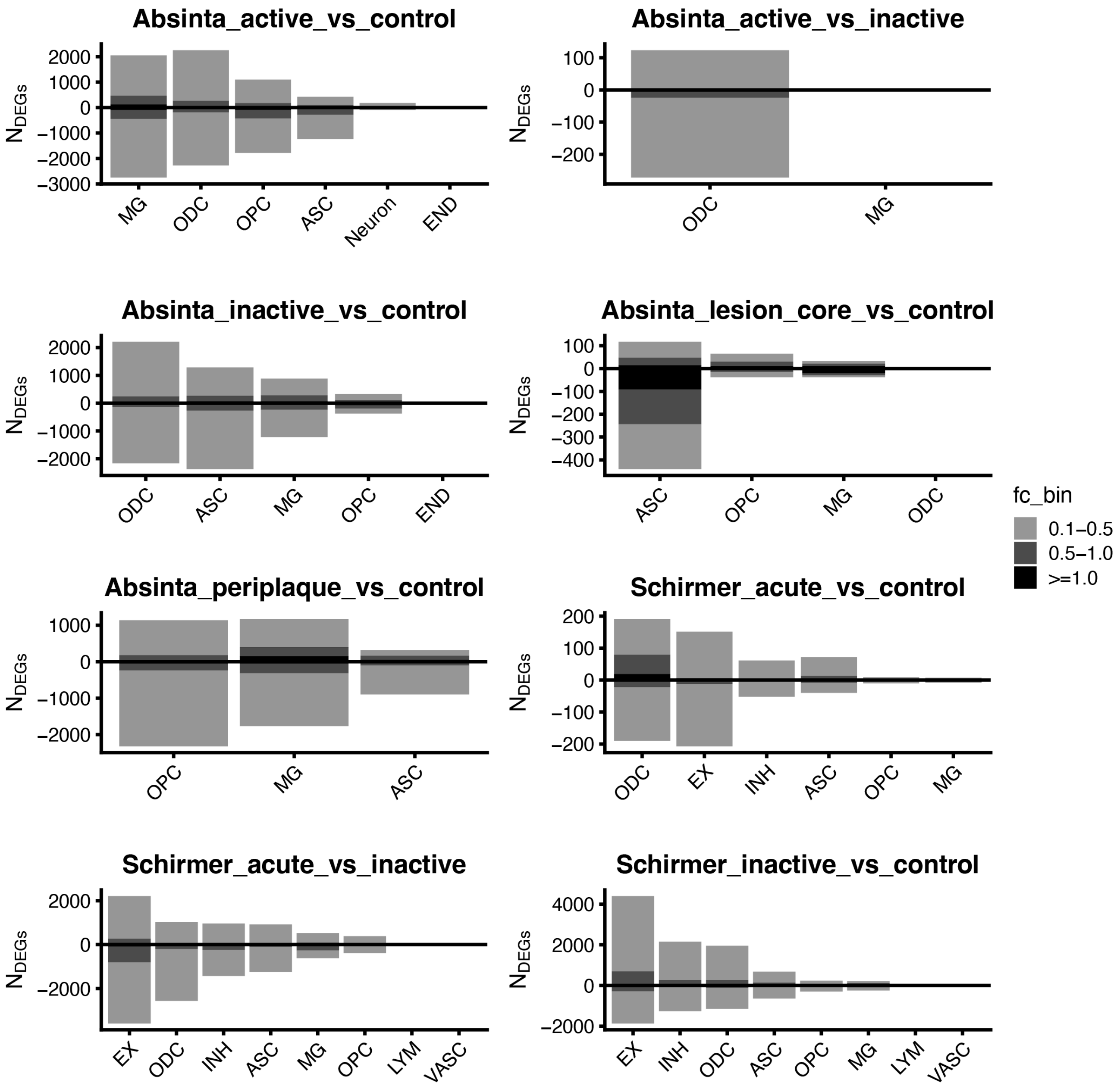


**Supplementary Fig. 10 | Summary of differential expression analysis in human multiple sclerosis snRNA-seq.** Bar plots showing the number of significantly differentially expressed genes in each comparison, across each major cell lineage in the human snRNA-seq datasets. In all differential expression tests, we compared disease vs. control, upregulated in the disease conditions corresponding to a positive fold change (above the Y=0 line in the bar graphs). Each bar is stratified by different $log_2$(fold change) bins.

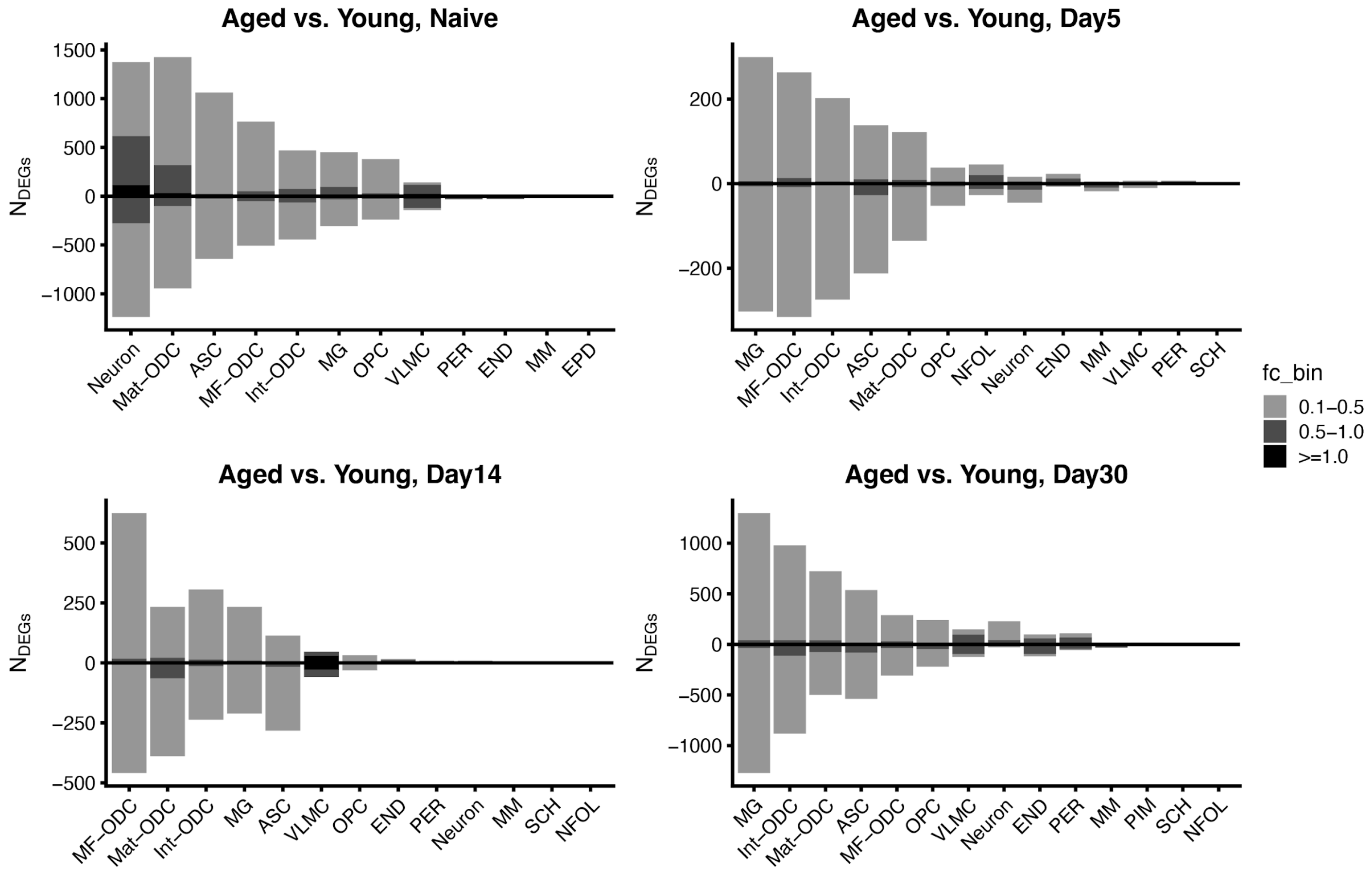


**Supplementary Fig. 11 | Summary of differential expression analysis in the mouse snRNA-seq dataset.** Bar plots showing the number of significantly differentially expressed genes in each comparison, across each major cell lineage in the mouse snRNA-seq dataset. Positive fold change (above the Y=0 line in the bar graphs) corresponds to DEGs upregulated in aged mice relative to young. Each bar is stratified by different $log_2$(fold change) bins.

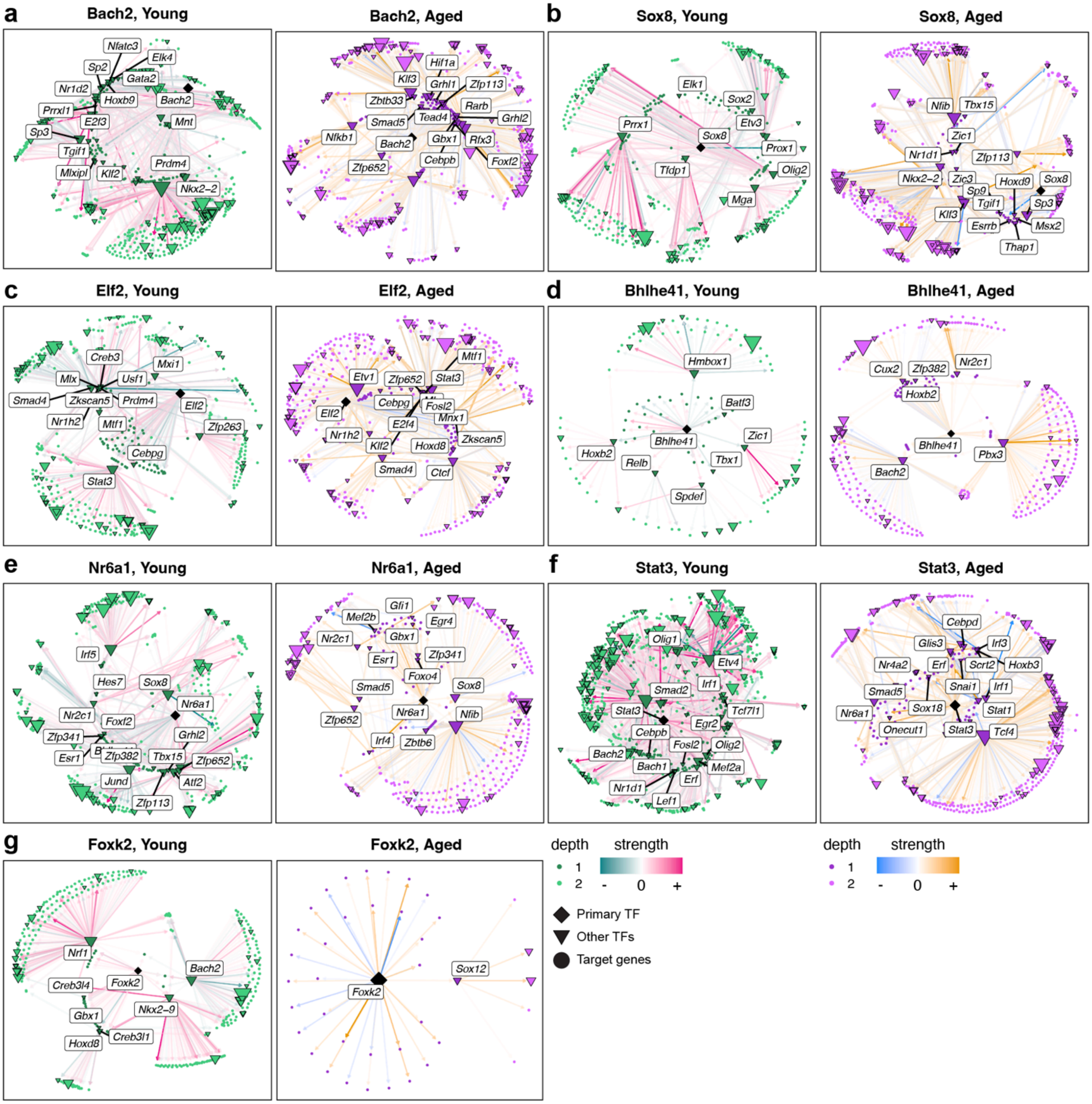


**Supplementary Fig. 12 | Transcription factor regulatory networks in TFs selected for downstream experiments.** Transcription factor (TF) regulatory network in young and aged mice for the seven TFs selected for downstream experiments: *Bach2* (**a**), *Sox8* (**b**), *Elf2* (**c**), *Bhlhe41* (**d**), *Nr6a1* (**e**), *Stat3* (**f**), and *Foxk2* (**g**). Nodes represent genes, and edges represent TF-gene regulatory relationships. Primary (direct) target genes representing the regulon of the main TF of interest (for example, *Bach2* in panel **a**) are shown as darker nodes, and secondary target genes (i.e., the regulons of other TFs targeted by the main TF) are shown in a lighter color. Other TFs targeted by the main TF are labeled.

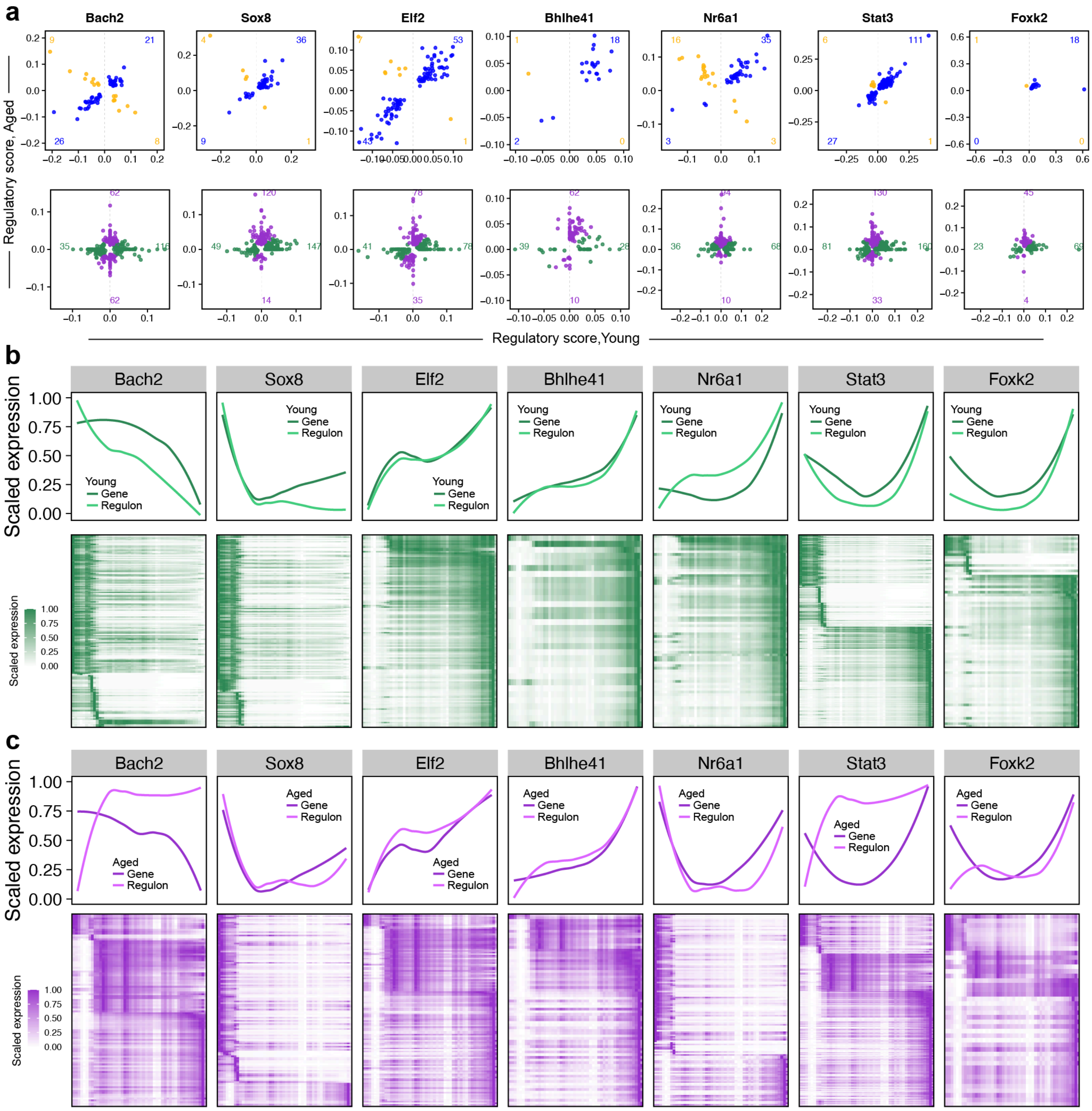


**Supplementary Fig. 13 | Transcription factor expression dynamics in TFs selected for downstream experiments. a**, Comparison of TF regulatory scores for the inferred regulons (target genes) of *Bach2, Sox8, Elf2, Bhlhe41, Nr6a1, Stat3,* and *Foxk2* in young vs. aged mice. Plots are faceted based on shared genes identified in the regulon in both young and aged (top) and unique genes (bottom). **b**, TF dynamics along the oligodendrocyte pseudotime trajectory in young mice. Top: scaled expression and aggregated TF regulon UCell[5] signature scores along the pseudotime trajectory for the same TFs shown in panel (**a**) Bottom: Heatmaps of scaled gene expression values for the inferred target genes of each TF in young mice. **c**, TF dynamics plots as shown in panel (**b**) for aged mice.

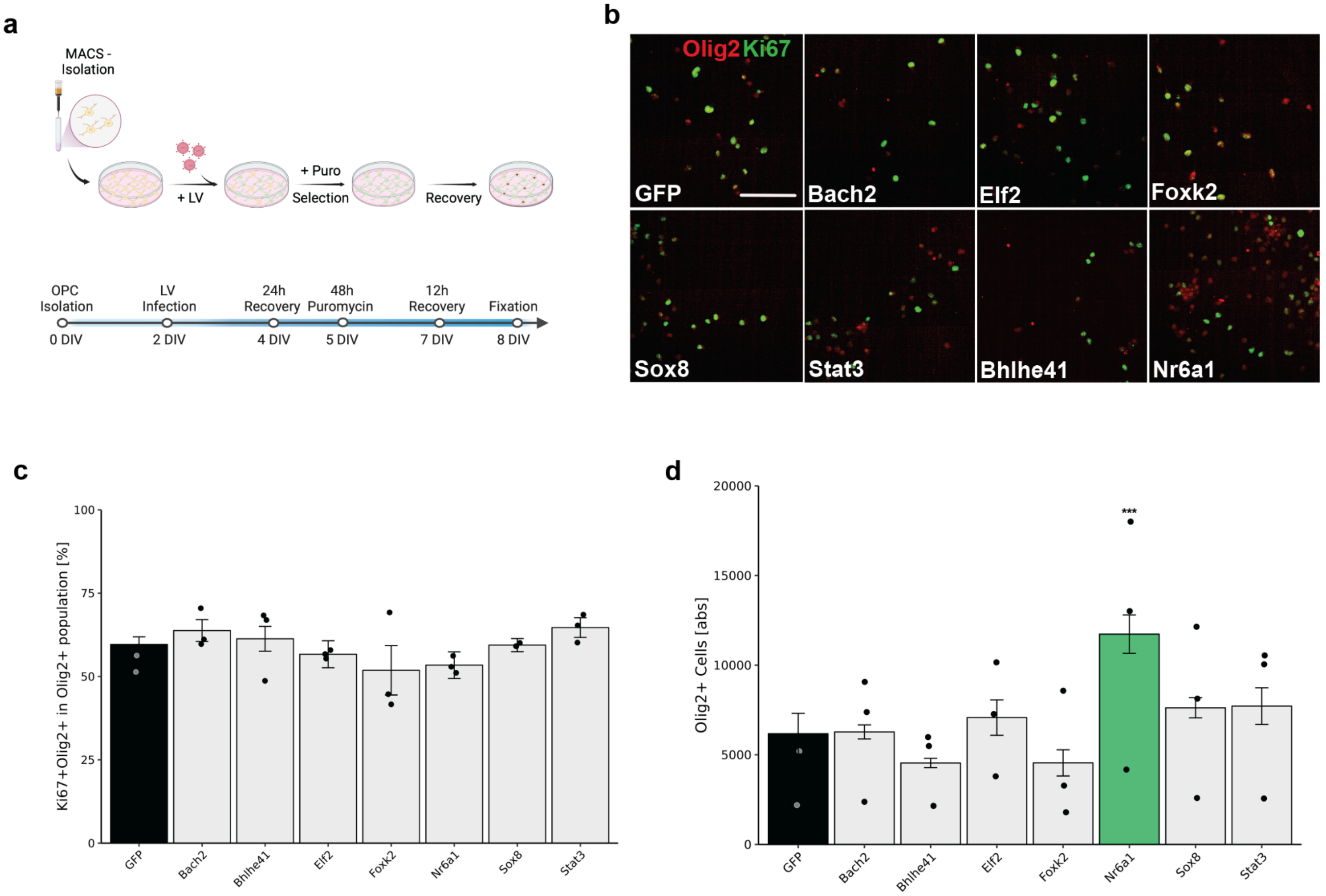


**Supplementary Fig. 14 | Effects of transcription factor overexpression on OPC proliferation and survival. a.** Diagram showing the experimental timeline of the OPC proliferation assay with lentiviral constructs for select transcription factor overexpression. **b.** Representative images of control OPCs (GFP) and OPCs transfected with the 7 select transcription factors in proliferation conditions and stained for Olig2 (red) and Ki67 (green). **c.** Quantification of the percentage of proliferating OPCs (Olig2+Ki67+) of total OLCs (Olig2+) with the different transcription factor overexpression treatments. Each data point shows the mean percentage across technical replicates (n=3) per biological replicate (n=3). Each bar shows the mean percentage across biological replicates (n=3), with propagated standard deviations indicated as error bars.**d.** Quantification of the absolute amount of total OLCs (Olig2+) with the different transcription factor overexpression treatments. Each data point shows the mean count across technical replicates (n=3) per biological replicate (n=3). Each bar shows the mean count across biological replicates (n=3), with propagated standard deviations indicated as error bars. Statistical analysis was performed using a linear mixed-effects model on the logit-transformed percentages (**c**) or the untransformed cell counts (**d**), with transcription factor overexpression as a fixed effect and animal as a random effect. Green bar highlights transcription factor overexpression treatment with a significant effect on the absolute number of olig2+ cells (**d**) compared to the control group (*** FDR-adjusted q < 0.001 ); treatments without a significant effect are indicated by white bars. Scale bar = 100 µm.

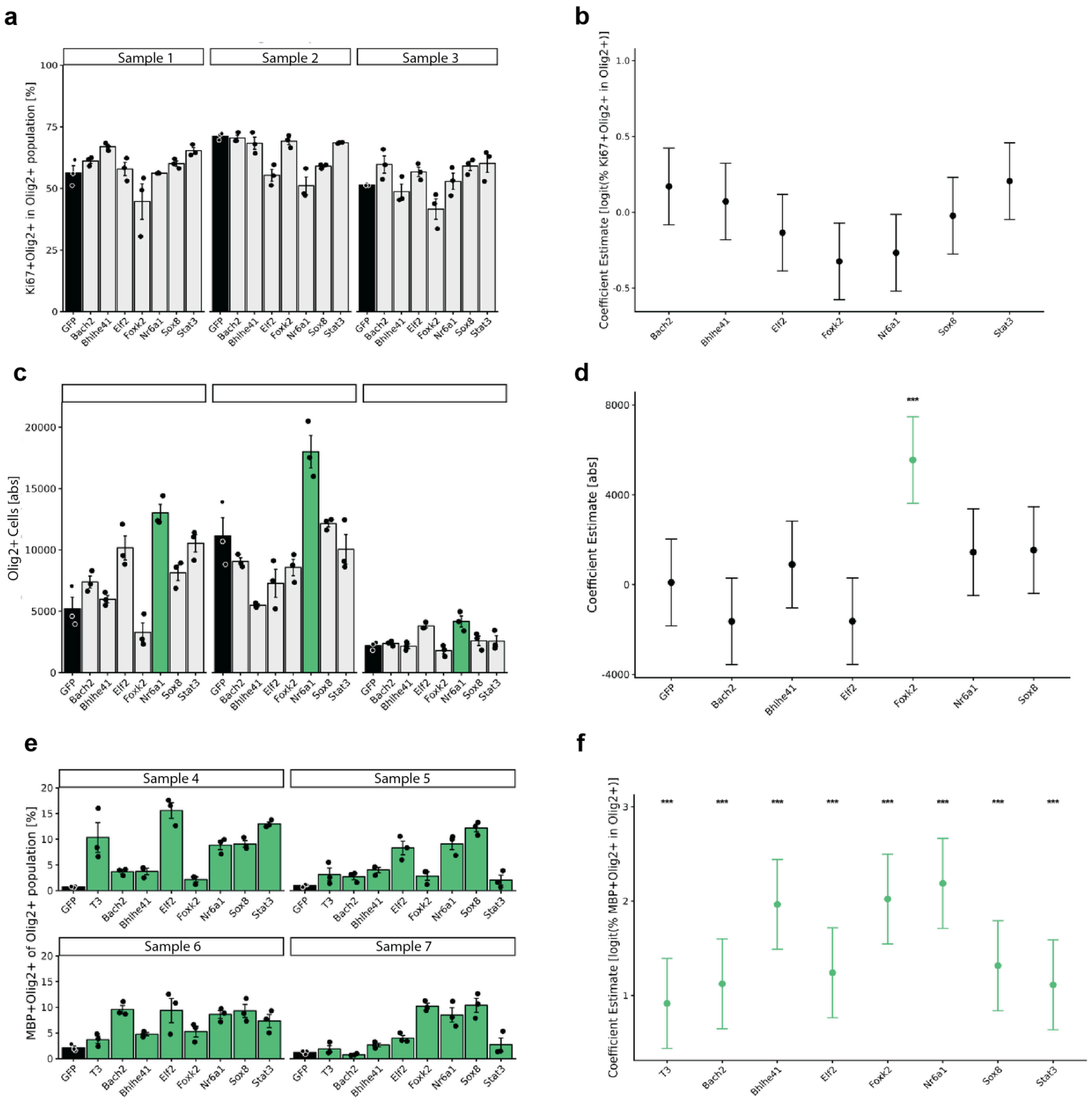


**Supplementary Fig. 15 | Effects of transcription factor overexpression on OPC proliferation, survival, and differentiation.** **a., c., e.** Quantification of the percentage of Ki67+ cells of total OLCs (Olig2+) (**a**), the absolute number of OLCs (**c**), and the percentage of differentiated OPCs (MBP+Olig2+) of total OLCs (Olig2+) (**e**), in response to transcription factor overexpression treatments. Each panel shows the mean value across technical replicates (n=3, standard errors indicated as error bars), faceted by biological sample. Green bars highlight transcription factor overexpression treatments with a significant effect on the percentage (**a**, **e**) or counts (**c**) compared to the control group (see Methods). **b., d., f.** Linear model coefficient estimates for the effect of transcription factor overexpression on the percentage of Ki67+ cells of total OLCs (Olig2+) (**b**), the absolute number of OLCs (**d**), and the percentage of differentiated OPCs (MBP+Olig2+) of total OLCs (Olig2+) (**f**) (see Methods). Scatter points indicate the coefficient estimates, with bars indicating the 95% confidence interval.

Green bars highlight transcription factor overexpression treatments with a significant effect on the percentage (**b**, **e**) or counts (**d**) compared to the control group (*** FDR-adjusted q < 0.001).

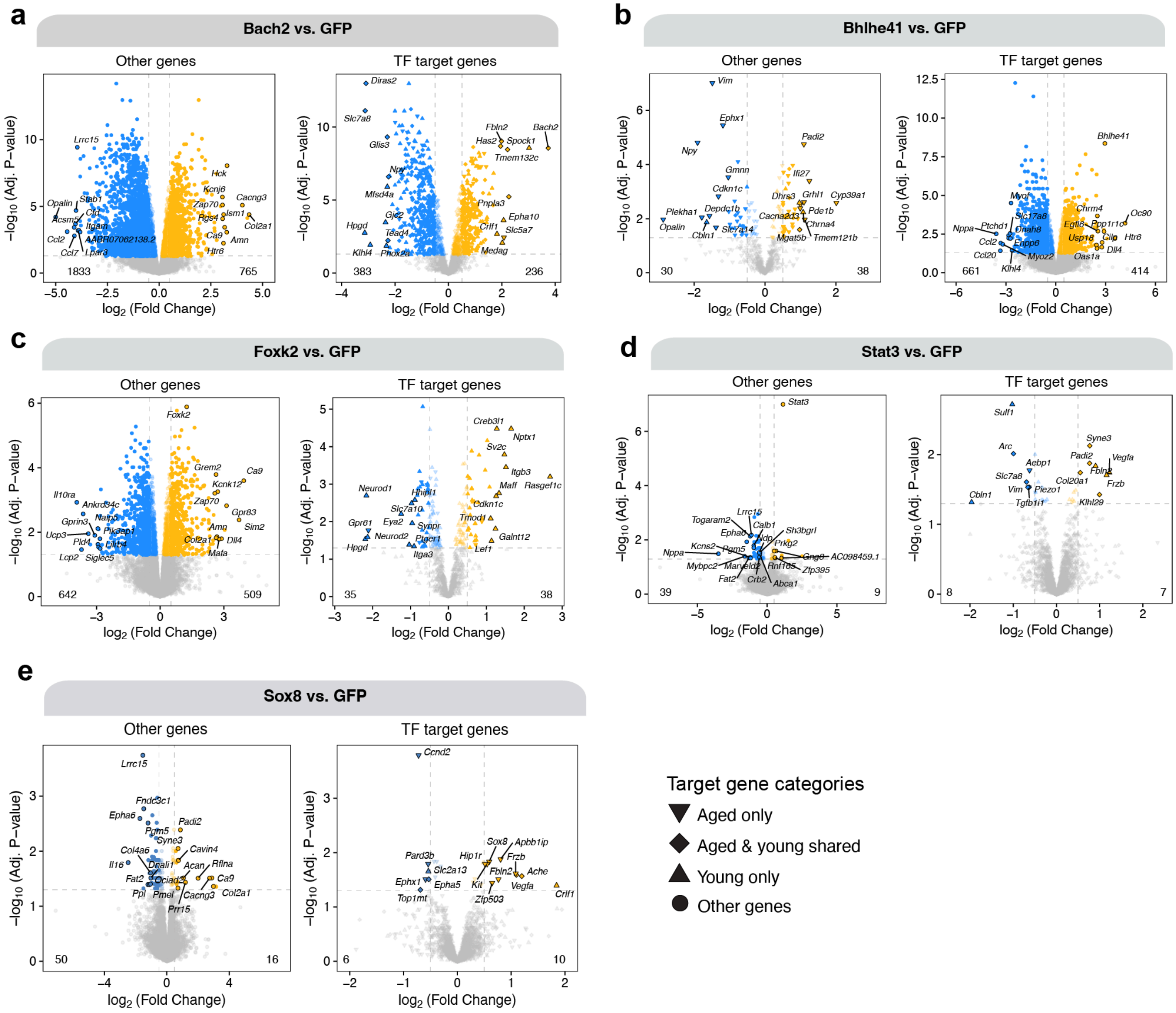


**Supplementary Fig. 16 | Differential expression results for bulk RNA-seq of transcription factor overexpression experiments.** Volcano plots showing differential gene expression results comparing transcription factor (TF) overexpression and GFP controls with bulk RNA-seq (n=4). The following TFs were overexpressed: *Bach2* (**a**), *Bhlhe41* (**b**), *Foxk2* (**c**), *Stat3* (**d**) and *Sox8* (**e**). For each panel, the plot on the right shows primary & secondary target genes from both young and aged mice for the TF of interest, and the plot on the left shows all other genes. Differential expression was determined by fitting a linear model to the expression data (see Methods), and genes were considered differentially expressed at a FDR-adjusted q-value < 0.05.
